# Large-scale genetic association and single cell accessible chromatin mapping defines cell type-specific mechanisms of type 1 diabetes risk

**DOI:** 10.1101/2021.01.13.426472

**Authors:** Joshua Chiou, Ryan J Geusz, Mei-Lin Okino, Jee Yun Han, Michael Miller, Paola Benaglio, Serina Huang, Katha Korgaonkar, Sandra Heller, Alexander Kleger, Sebastian Preissl, David U Gorkin, Maike Sander, Kyle J Gaulton

## Abstract

Translating genome-wide association studies (GWAS) of complex disease into mechanistic insight requires a comprehensive understanding of risk variant effects on disease-relevant cell types. To uncover cell type-specific mechanisms of type 1 diabetes (T1D) risk, we combined genetic association mapping and single cell epigenomics. We performed the largest to-date GWAS of T1D in 489,679 samples imputed into 59.2M variants, which identified 74 novel association signals including several large-effect rare variants. Fine-mapping of 141 total signals substantially improved resolution of causal variant credible sets, which primarily mapped to non-coding sequence. To annotate cell type-specific regulatory mechanisms of T1D risk variants, we mapped 448,142 candidate *cis-*regulatory elements (cCREs) in pancreas and peripheral blood mononuclear cell types using snATAC-seq of 131,554 nuclei. T1D risk variants were enriched in cCREs active in CD4+ T cells as well as several additional cell types including pancreatic exocrine acinar and ductal cells. High-probability T1D risk variants at multiple signals mapped to exocrine-specific cCREs including novel loci near *CEL, GP2* and *CFTR*. At the *CFTR* locus, the likely causal variant rs7795896 mapped in a ductal-specific distal cCRE which regulated *CFTR* and the risk allele reduced transcription factor binding, enhancer activity and *CFTR* expression in ductal cells. These findings support a role for the exocrine pancreas in T1D pathogenesis and highlight the power of combining large-scale GWAS and single cell epigenomics to provide insight into the cellular origins of complex disease.

## INTRODUCTION

Type 1 diabetes (T1D) is a complex autoimmune disease characterized by the loss of insulin-producing pancreatic beta cells and subsequent hyperglycemia^1^, where the triggers of autoimmunity and disease onset remain poorly understood. T1D has a strong genetic component, most prominently at the major histocompatibility complex (MHC) locus but including 60 additional risk loci identified in genome-wide and targeted array association studies^2–6^. T1D associated variants at risk loci are largely non-coding, and intersection of T1D associated variants with epigenomic data has identified an enrichment of risk variants within lymphoid enhancers^2^. However, due to limited sample sizes, incomplete variant coverage, and the limited cell type resolution of existing epigenomic maps, the causal variants and cellular mechanisms of action of T1D risk loci are largely unresolved.

## RESULTS

### Comprehensive discovery and fine mapping of T1D risk signals

To discover novel risk loci and improve fine mapping of causal variants for T1D, we performed a genome-wide association study (GWAS) of 18,803 T1D cases and 470,876 controls of European ancestry from 9 country-of-origin and array-matched cohorts (**Supplemental Table 1**). After applying uniform quality-control measures (**Supplemental Figure 1**), where we removed low-quality genotypes, individuals of non-European ancestry, or controls with other autoimmune diseases, we imputed genotypes into the TOPMed r2 panel and tested for T1D association^7^. Through meta-analysis, we combined association results for 59,244,856 variants across cohorts and observed 80 loci reaching genome-wide significance (P<5×10^−8^), including 30 loci previously unreported in T1D risk (**Figure 1a, Supplemental Figure 2, Supplemental Table 2**). Previous studies have identified independent association signals at multiple T1D loci^2^, and we reasoned that our increased sample size would uncover additional independent signals. Through iterative conditional analyses, we discovered 52 secondary signals at locus-wide significance (P<1×10^−5^), of which 44 were previously unknown (**Supplemental Figure 3, Supplemental Table 2**). Over 40% (36/89) of loci contained more than one independent signal; for example, the known *BACH2* locus and novel *BCL11A* locus each had three signals (**Figure 1b**), and at the *IL2RA* locus we identified six independent signals, three of which were novel (**Supplemental Figure 3**).

**Fig 1.**
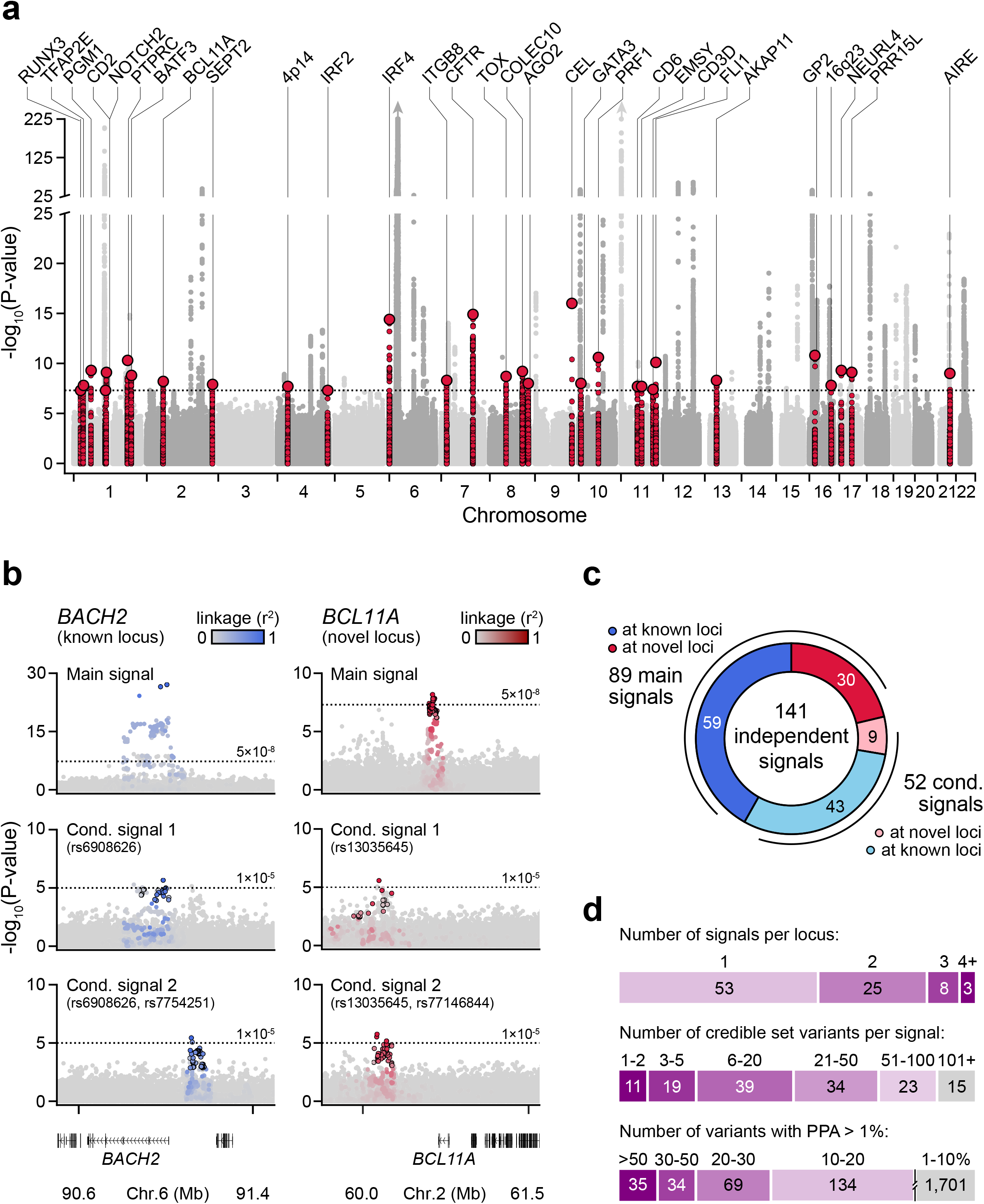
Genome-wide association and fine mapping identifies novel signals for T1D risk. (a) Manhattan plot showing genome-wide T1D association p-values (-log10 transformed). Novel loci are colored in red and labeled based on the nearest gene, and index variants have larger radii and are circled. The dotted line indicates genome-wide significance (P=5×10^−8^). (b) Locus plots showing independent association signals at the known *BACH2* locus (left) and the novel *BCL11A* locus (right). For conditional signals, the variants used for conditional analysis are indicated under the title in parentheses. Variants are colored (known=blue, novel=red) based on linkage disequilibrium (r^2^) with the index variant for each signal. The dotted line indicates the genome-wide significance threshold (P=5×10^−8^) for the main signal and the locus wide significance threshold (P=1×10^−5^) for the conditional signals. (c) Breakdown of 141 independent T1D risk signals after conditional fine-mapping analyses. Among these were 89 main signals at 59 known loci (excluding the MHC region) and 30 novel loci, and 52 conditional signals including 43 at known loci and 9 at novel loci. (d) Breakdown of the number of signals per locus (top), number of 99% credible set variants per signal from fine mapping (middle), and the number of variants with posterior probability of association >1% (bottom).

**Fig 2.**
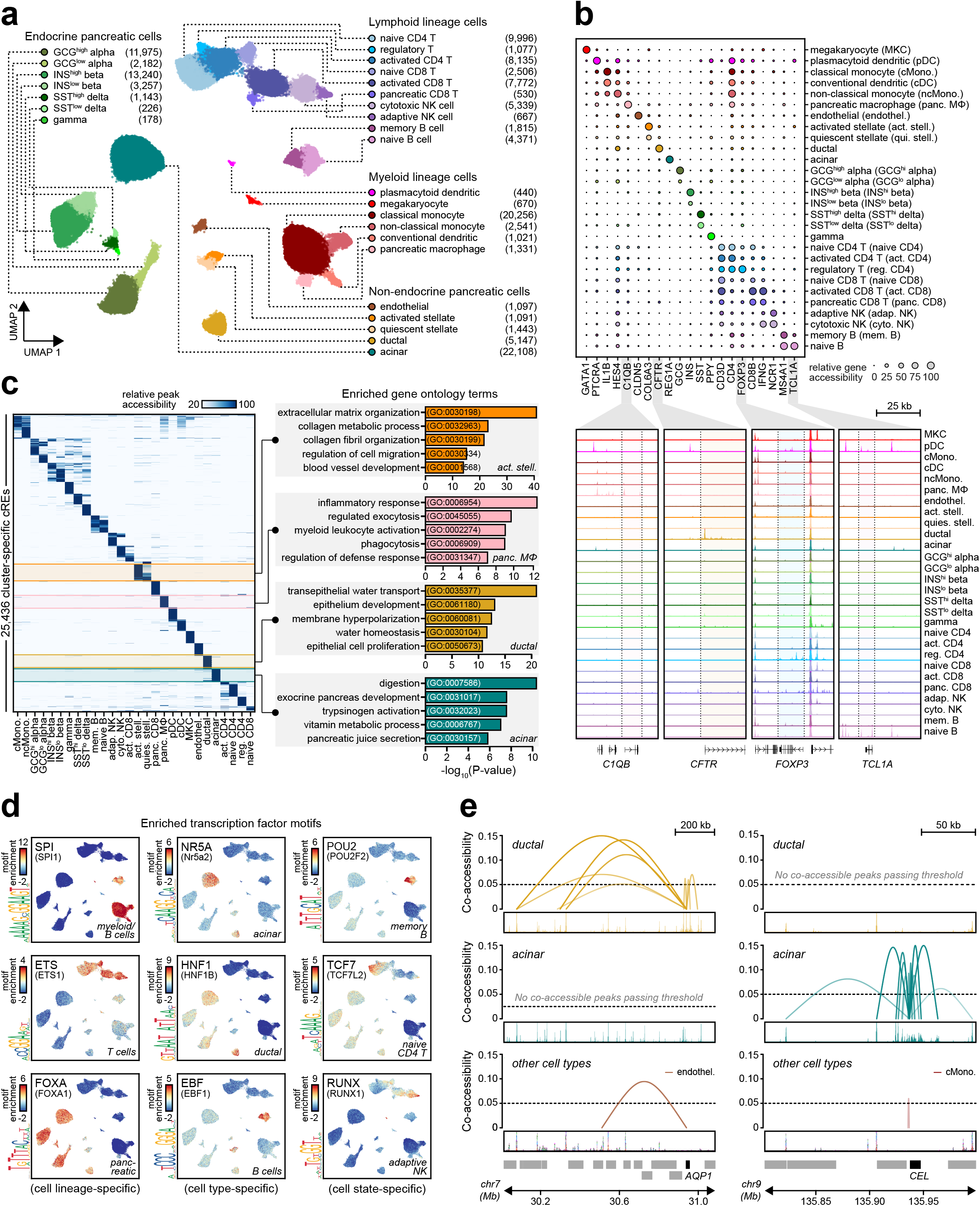
Comprehensive reference map of 131,554 single cell chromatin accessibility profiles from T1D-relevant tissues. (a) Clustering of accessible chromatin profiles from 131,554 cells from single cell experiments of peripheral blood mononuclear cells, whole pancreas tissue, and purified pancreatic islets. Cells are plotted on the first two UMAP components and colored based on cluster assignment. Clusters are grouped into categories of cell types, and the number of cells in each cluster are shown next to its corresponding label. (b) Dot plot (top) of relative gene accessibility (chromatin accessibility reads across gene bodies, averages for each cluster and scaled from 0-100 across columns/clusters) showing examples of marker genes used to identify cluster labels. Circle sizes are scaled according to the relative gene accessibility value. Genome browser tracks (bottom) showing aggregated chromatin accessibility profiles in a 50 kb window around selected marker genes. (c) Relative peak accessibility for 25,436 cluster-specific peaks across all 28 clusters (left), and enriched gene ontology terms with GREAT for peaks specific to pancreatic macrophages, activated stellate, ductal, and acinar cells (right). (d) Single cell motif enrichment z-scores for TFs showing specificity for cell lineage (SPI – myeloid and B cells, ETS – T cells, FOXA – pancreatic), cell type (NR5A – acinar, HNF1 – ductal, EBF – B cells), and cell state (POU2 – memory B, TCF7 – naïve CD4 T, RUNX – adaptive NK). The sequence logo for the enriched motif is displayed to the left of each UMAP plot. (e) Examples of cell type-specific co-accessibility between the promoter of AQP1 and distal sites in ductal cells (left, chr7:30,000,000-31,100,000, scale: 0-10 CPM) and the promoter of CEL and distal sites in acinar cells (right, chr9:135,800,000-136,000,000, scale: 0-10 CPM).

**Fig 3.**
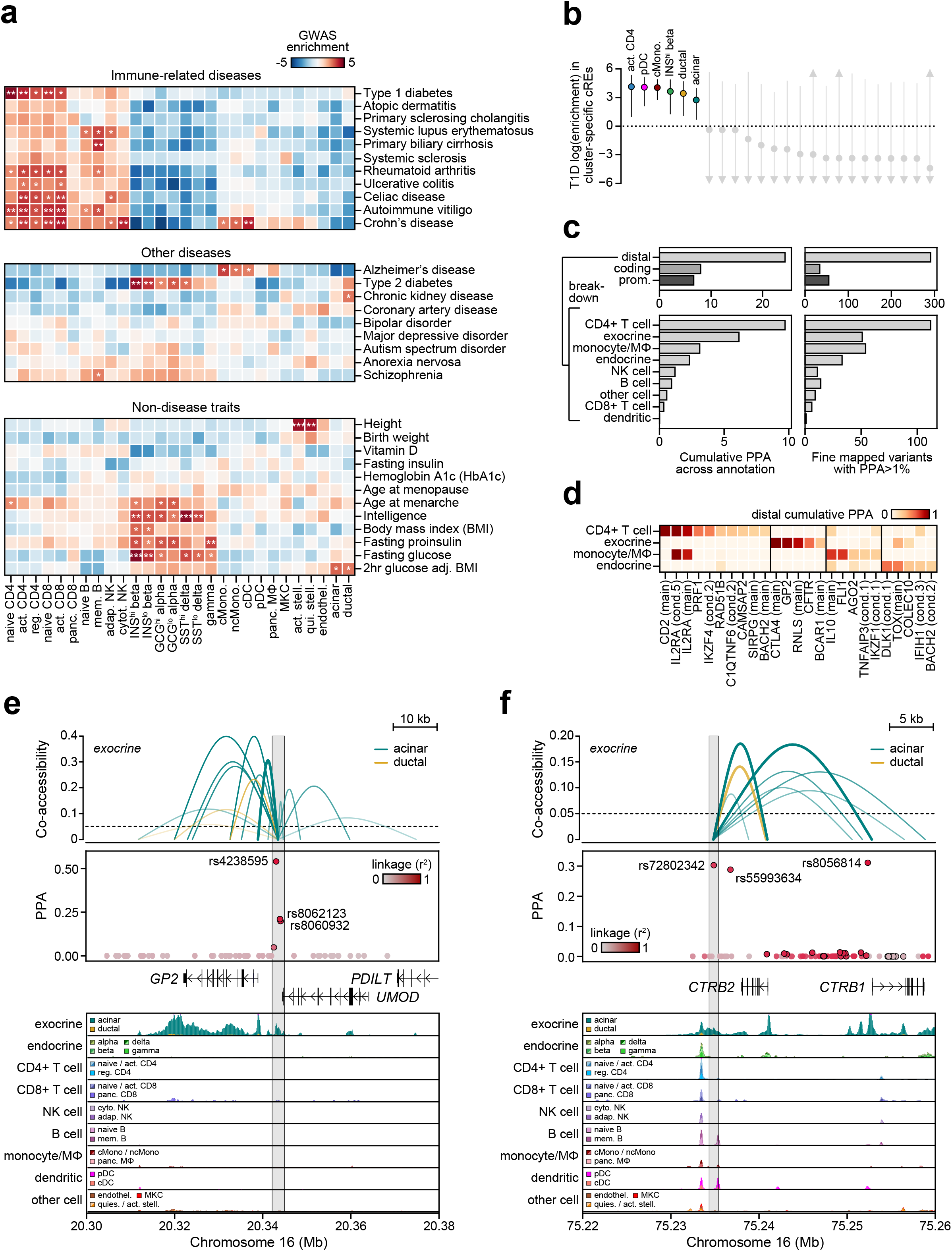
Cell type-specific enrichment and mechanisms of T1D risk variants. (a) Relative LD score regression enrichment z-scores (enrichment relative to background genomic annotations including a merged set of all peaks) for autoimmune and inflammatory diseases (top), other diseases (middle), and non-disease quantitative endophenotypes (bottom) for cCREs active in pancreatic and blood cell types and states. ***FDR<.001 **FDR<.01 *FDR<.1. (b) T1D enrichment within cell type-specific cCREs. Labeled clusters have a positive enrichment estimate. Points represent log-transformed fgwas enrichment estimates and lines represent 95% confidence intervals. (c) Breakdown of cumulative fine mapping probability (PPA) (left) and fine mapped variants (right). Variants and their probabilities are assigned without replacement to annotations from top to bottom. Variants are first broken down by genomic annotations (top), and variants overlapping a distal peak are further broken down by cell type groups (bottom). CD4 T cell: naïve CD4 T + activated CD4 + regulatory T; exocrine: acinar + ductal; endocrine: GCG^high^ alpha + GCG^low^ alpha + INS^high^ beta + INS^low^ beta + SST^high^ delta + SST^low^ delta + gamma; monocyte/MΦ: classical monocyte + non-classical monocyte + pancreatic macrophage; NK cell: cytotoxic NK + adaptive NK; B cell: naïve B + memory B; CD8 T cell: naïve CD8 T + activated CD8 T + pancreatic CD8 T; other cell: megakaryocytes + activated stellate + quiescent stellate + endothelial; dendritic: conventional dendritic + plasmacytoid dendritic. (d) Signals with the highest cumulative PPA for cell type groups with at least 2.5 cumulative PPA. (e) The *GP2* signal contains 3 variants (rs4238595, rs8060932, and rs8060932) in a distal peak upstream of the *GP2* promoter (top, chr16:20,300,000-20,380,000). These variants are linked to *GP2* through co-accessibility in acinar cells and account for the majority of the causal probability (cumulative PPA=.98) for the signal (middle). Genome browser tracks (bottom) show that chromatin accessibility at both the peak and the *GP2* promoter is highly specific to acinar cells. (f) The top variant at the *CTRB1/2/BCAR1* signal rs72802342 (middle) overlaps a distal peak co-accessible with the *CTRB2 and CTRB1* promoters in acinar cells (top: chr16:75,220,000-75,260,000, hg19). Genome browser tracks (bottom, scale: 0-15) show that chromatin accessibility at the *CTRB1* and *CTRB2* promoters are highly specific to acinar cells. Fine mapped variants are colored based on linkage disequilibrium to the index variant. Variants contained in the 99% credible set are circled in black.

The TOPMed r2 panel enables more accurate imputation of rare variants over previous reference panels, and in our study, we identified five novel T1D-associated variants with minor allele frequency (MAF) less than 0.005 and large effects on disease risk (**Supplemental Table 2, Supplemental Figure 4**). Among these rare variants, rs541856133 (MAF=.0015, OR=2.97) mapped to a non-coding region directly upstream of *CEL*, which has been implicated previously as the cause of maturity-onset diabetes of the young with pancreatic exocrine dysfunction (MODY8)^8^. We also identified a novel protein-coding protective variant at *IFIH1* (p.Asn160Asp, rs75671397, MAF=.002, OR=0.32), which was conditionally independent of the known protein-coding variant signals in this gene. The three additional rare T1D risk variants mapped to non-coding regions at the 16q23 (rs138099003, MAF=.0015, OR=2.29), *SH2B*3 (rs762349492, MAF=.0018, OR=1.99), and *TOX* (rs192456638, MAF=.0045, OR=1.80) loci (**Supplemental Table 2, Supplemental Figure 4**).

**Fig 4.**
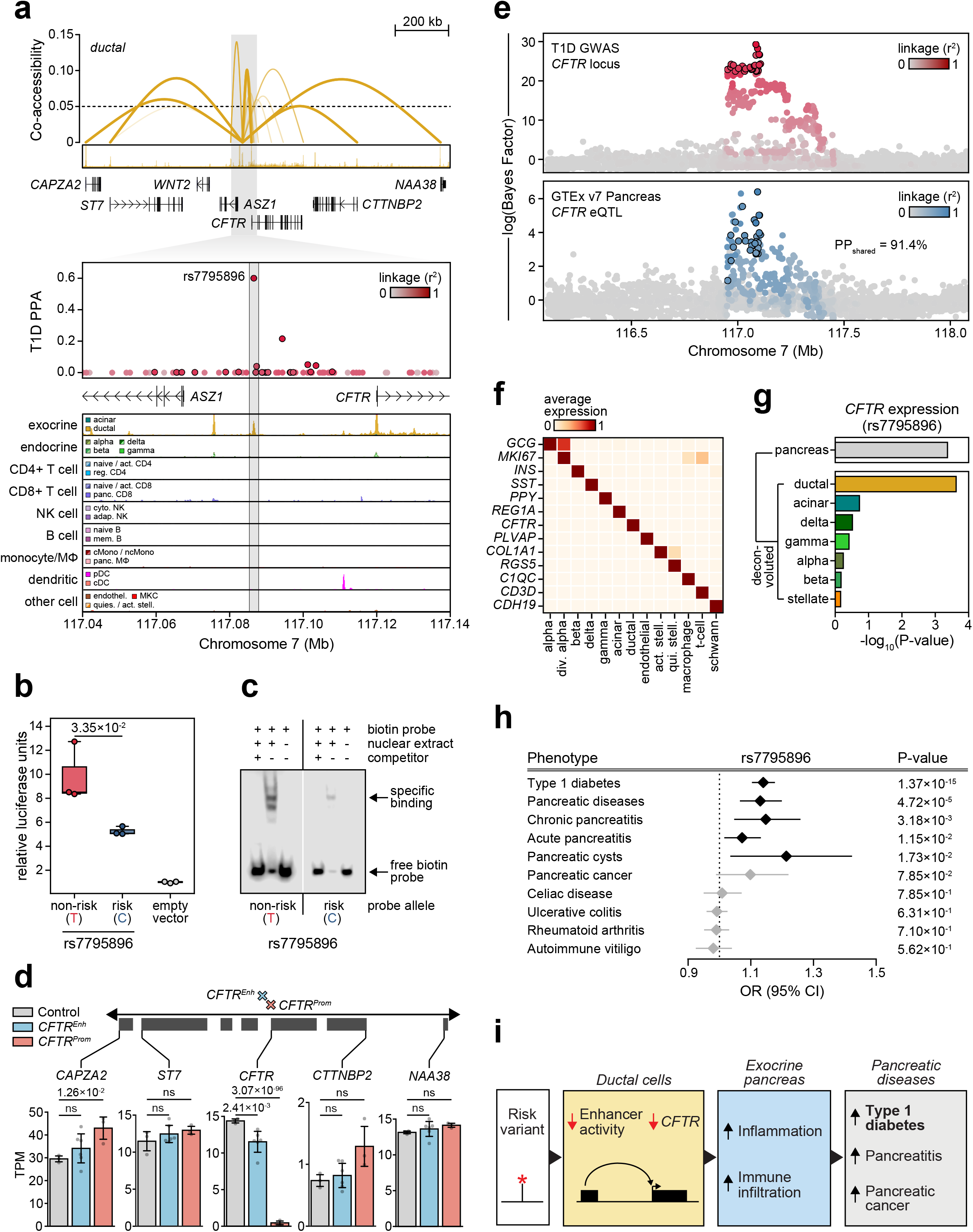
Fine-mapped variant at the *CFTR* locus mediates T1D risk through distal regulation of *CFTR* in pancreatic ductal cells. (a) The *CFTR* locus contains a single fine-mapped variant (rs7795896) in a distal cCRE linked to the promoter of *CFTR* and several other genes through co-accessibility (top; region shown: chr7:116,490,000-117,860,000). The cCRE is located approximately 33 kb upstream of the *CFTR* promoter. Zoomed-in view (chr7:117,040,000-117,140,000, scale: 0-5 CPM) of fine mapped variants (middle) and genome browser tracks (bottom) at this locus show that the cCRE is highly specific to ductal cells. (b) Luciferase reporter assay in Capan-1 cells transfected with pGL4.23 minimal promoter plasmids containing rs7795896 in the forward orientation. Relative luciferase units represent Firefly:Renilla ratios normalized to control cells transfected with the empty vector. P-values are from a two-tailed Student’s t-test. (c) Electrophoretic mobility shift assay (EMSA) with nuclear extract from Capan-1 cells using probes from both alleles of rs7795896. Bands with specific binding are labeled. (d) CRISPR interference-mediated inactivation of the distal site containing rs7795896 (*CFTR*^iEnh^; 2 guide RNAs; 3 replicates; n=6 total) or the *CFTR* promoter (*CFTR*^iProm^; n=3 replicates) in CAPAN-1 cells. Differential analysis of genes with promoters co-accessible with the peak show that CFTR expression is significantly reduced in both *CFTR*^iProm^ and *CFTR*^iEnh^ cells. Data are shown as transcripts per million (TPM). Error bars show 95% confidence interval and datapoints underlying each boxplot are shown. (e) Bayesian colocalization showing that the T1D risk signal (top) and *CFTR* pancreas eQTL from GTEx v7 (bottom) are likely driven by the same causal variant. Variants are colored based on the linkage disequilibrium to the index variant. Variants in the 99% credible set are circled in black. (f) Heatmap showing the average expression (normalized counts, scaled from 0-1 across cell types) of marker genes of different pancreatic cell types from single cell RNA-seq. *CFTR* expression is highly specific to ductal cells. (g) Deconvolution of the *CFTR* pancreas eQTL using *in-silico* cell type proportion estimation and re-analyses of GTEx pancreas data using interaction analyses shows that the eQTL signal only has a significant interaction with ductal cell proportion. (h) Forest plot showing association of pancreatic disease traits in a meta-analysis of UK Biobank and FinnGen data for rs7795896 compared to association of autoimmune traits from large European GWAS. (i) Variants regulating genes with specialized function in the exocrine pancreas influence risk of type 1 diabetes. At the *CFTR* locus, a variant reducing ductal cell enhancer activity and *CFTR* expression increases risk of T1D and other pancreatic disease, and we hypothesize that these effects are mediated through inflammation and immune infiltration in the exocrine pancreas.

We next sought to fine map causal variants of T1D signals using a Bayesian approach^9^. In total we considered 141 signals including 89 primary and 52 conditional signals at known and novel loci excluding the MHC locus due to complex LD structure (**Figure 1c**). We defined linkage disequilibrium (LD)-based credible sets for the 141 signals, using new index variants at known loci where applicable. For each signal, we then used approximate Bayes factors^9^ to calculate the posterior probability of association (PPA) for each variant and defined credible sets of variants that summed up to 99% cumulative PPA (**Supplemental Table 3**). Compared to previous efforts^2,10^, our fine-mapping resolution was drastically improved based on two complementary measures: 1) fewer number of credible set variants per signal (median 24 variants) and 2) a greater number of variants with high causal probabilities (**Figure 1d**). At nearly half of all T1D signals (49%; 69/141) the credible set contained 20 or fewer variants, and 25% (35/141) contained a single variant explaining the majority of the posterior probability (>50% PPA). Among credible set variants, 23 variants with PPA>1% were nonsynonymous changes, including several at novel loci p.Arg471Cys in *AIRE* (PPA=.99), p.Val11Ile in *BATF3* (PPA=.081), p.Ala91Val in *PRF1* (PPA=0.038), and p.Val131Phe in *CD3G* (PPA=.028) (**Supplemental Table 4**).

Given our comprehensive genome-wide T1D genetic association and fine-mapping data, we used these data to derive insight into disease pathophysiology. We therefore broadly characterized relationships between T1D and other complex traits and diseases by performing genome-wide genetic correlation analyses using LD score regression. As expected, T1D had significant (FDR<.10) positive correlations with autoimmune diseases including rheumatoid arthritis (r_g_=0.43, FDR=7.34×10^−5^), systemic lupus erythematosus (r_g_=0.36, FDR=2.52×10^−7^), celiac disease (r_g_=0.28, FDR=1.11×10^−3^), and autoimmune vitiligo (r_g_=0.30, FDR=2.02×10^−5^), as well as a negative correlation with ulcerative colitis (r_g_=-0.17, FDR=2.94×10^−3^) (**Supplemental Figure 5**). Among other traits, we observed significant positive correlations with metabolic traits and diseases such as fasting proinsulin (r_g_=0.18, FDR=8.91×10^−2^) and fasting insulin level, (r_g_=0.18, FDR=6.85×10^−3^), coronary artery disease (r_g_=0.12, FDR=6.85×10^−3^) and type 2 diabetes (r_g_=0.10, FDR=4.39×10^−3^), and positive correlations with pancreatic diseases such as pancreatic cancer (r_g_=0.25, FDR=7.40×10^−2^) and chronic pancreatitis (r_g_=0.13, FDR=3.84×10^−1^), although the latter estimate was not significant. These results demonstrate relationships between genetic effects on T1D risk and a diversity of traits including autoimmune, pancreatic and metabolic disease.

### Defining cell type-specific *cis*-regulatory programs in T1D-relevant tissues

The large majority of T1D risk signals map to non-coding regions and likely affect gene regulation^2^. In order to annotate gene regulatory programs affected by T1D risk variants, we generated a reference map of cell type-specific accessible chromatin using single nucleus ATAC-seq (snATAC-seq) assays of T1D-relevant tissues including peripheral mononuclear blood cells (PBMC), purified pancreatic islets, and whole pancreas tissue from non-diabetic donors (**Supplemental Table 5**). To cluster cells obtained from these assays, we used a modified version of our previous pipeline^11^ that included rigorous quality control, removal of potential doublets, and removal of potential confounding effects between different donors, tissues, and technologies to group 131,554 chromatin accessibility profiles into 28 clusters (**Figure 2a, Supplemental Figure 6**). We assigned cell type identity to each cluster using the chromatin accessibility profiles of gene bodies for known marker genes, and identified cells representing lymphoid, myeloid, endocrine, exocrine, endothelial, and stellate cell types (**Figure 2a-b**). Within lymphoid and myeloid cells, there were clusters representing both peripheral blood cells as well as tissue resident cells in the pancreas based on both marker gene accessibility and tissue-of-origin profiles (**Figure 2a-b, Supplemental Figure 6**). For example, we observed accessibility at *C1QB* marking pancreatic tissue-resident macrophages, at *REG1A* marking pancreatic acinar cells, and at *CFTR* marking pancreatic ductal cells (**Figure 2b**). We also observed distinct patterns of chromatin accessibility at marker genes between different clusters of the same cell type allowing us to further discriminate specific sub-types such as *FOXP3* for regulatory T cells relative to other T cells and *TCL1A* for naïve B cells relative to memory B cells (**Figure 2b**).

To characterize the regulatory programs of each cell type and cell state, we aggregated reads from cells within each cluster and called accessible chromatin sites representing candidate *cis-*regulatory elements (cCREs). Across all 28 clusters, we identified a total of 448,142 cCREs and an average of 77,812 cCREs per cluster (**Supplementary Data 1**). To further define regulatory programs defining the identity of each cell type, we calculated the relative accessibility of each cCRE across all clusters and identified 25,436 cell type-specific cCREs with accessibility patterns specific to a given cluster **(Figure 2c, Supplementary Data 2**). To confirm that cell type-specific cCREs regulated key processes involved in cellular identity, we identified gene ontology (GO) terms enriched for each set of cell type-specific cCREs using GREAT^12^. GO terms significantly enriched in cell type-specific cCREs represented highly specialized cellular processes, for example inflammatory response for pancreatic tissue-resident macrophages (P=6.09×10^−12^), extracellular matrix organization for activated stellate cells (P=1.47×10^−41^), transepithelial water transport for ductal cells (P=1.26×10^−21^) and digestion for acinar cells (P=1.18×10^−11^) (**Figure 2c, Supplementary Table 6**).

We next decoded the regulatory logic underlying cCRE activity for each cell type. First, we identified candidate transcription factors (TFs) regulating cCRE activity by identifying sequence motifs enriched in accessible chromatin of each cell type using chromVAR^13^. There were 290 motifs in JASPAR^14^ with evidence for variable enrichment across cell types (**Supplementary Table 7**). Enriched motifs included TF families with lineage-specific enrichment such as SPI in myeloid and B cells, ETS in T cells, and FOXA in pancreatic endocrine and exocrine cells^15–17^ (**Figure 2d**). We also identified motifs enriched in specific cell types such as NR5A in acinar cells^18^, HNF1 in ductal cells^19^, and EBF in B cells^20^ (**Figure 2d**), as well as motifs for TF families enriched in specific states within a cell type, such as POU2 in memory B cells^21^, TCF7 in naïve CD4+ T cells^22^, and RUNX in adaptive NK cells^23^ (**Figure 2d**). Second, we defined cell type-resolved links between distal cCREs and putative target gene promoters using co-accessibility across single cells with Cicero^24^. Considering all cell types, we observed a total of 1,028,428 links between distal cCREs and gene promoters (**Supplemental Data 3**), where 145,138 distinct distal cCREs were linked to at least one promoter. In many cases, co-accessible links were highly cell type-specific; for example, multiple distal cCREs were co-accessible with the *AQP1* promoter in ductal cells and the *CEL* promoter in acinar cells, none of which were identified in other cell types (**Figure 2e**). Together these results identify candidate transcriptional regulators and target genes of distal cCREs in pancreatic and immune cell types.

### Annotating fine-mapped T1D risk variants with cell type-specific regulatory programs

We reasoned that our cell type-resolved regulatory maps would enable deeper insight into pancreatic and blood cell types involved in T1D pathogenesis. We therefore determined enrichment of variants associated with T1D as well as other complex diseases^25–42^ and qualitative endophenotypes^43–52^ for cCREs using stratified LD score regression^53^. For T1D, the most significant enrichment was for variants in CD4+ T cell cCREs (naïve CD4+ T Z=4.54, FDR=1.26×10^−3^; activated CD4+ T Z=3.83, FDR=5.88×10^−3^; regulatory T Z=3.26, FDR=1.35×10^−2^) (**Figure 3a**). Notably, we did not observe evidence for enrichment in resident immune cells in the pancreas (pancreatic CD8+ T cell Z=0.46, FDR=0.93; pancreatic tissue-resident macrophage Z=-1.02, FDR=1.0). Outside of immune cell types, pancreatic ductal cell cCREs had the strongest T1D enrichment, although this estimate was not significant (ductal Z=0.46, FDR=0.93). Other immune-related diseases were also enriched within lymphocyte cCREs, although Crohn’s disease was also enriched for monocytes and conventional dendritic cell cCREs (**Figure 3a**). As expected, type 2 diabetes and glycemic traits were strongly enriched in pancreatic endocrine cell cCREs, but interestingly, glycemic traits such as glucose levels at 2 hours post-OGTT were also enriched in pancreatic acinar and ductal cell cCREs (**Figure 3a**). Together these results demonstrate that T1D associated variants are broadly enriched for CD4+ T cell cCREs, and highlight other complex traits and diseases enriched for pancreatic and immune cell type cCREs.

Despite the strong enrichment of T1D-associated variants in CD4+ T cells, less than half of fine-mapped T1D signals overlapped a CD4+ T cell cRE, suggesting that additional cell types contribute to T1D risk. In order to identify additional disease-relevant cell types, we used an orthogonal approach to test for enrichment of T1D variants within the subset of cCREs specific to each cell type (from **Figure 2c; see Methods**). As expected, T1D variants genome-wide were enriched in cCREs specific to CD4+ T cells (activated CD4+ T log enrich=4.14, 95% CI=0.97-5.37) as well as pancreatic beta cells (log enrich=3.64, 95% CI=1.23-4.90) (**Figure 3b**). Interestingly, T1D variants were also enriched in cCREs specific to plasmacytoid dendritic cells (log enrich=4.08, 95% CI=2.09-5.16), classical monocytes (log enrich=4.04, 95% CI=2.74-4.92), and pancreatic acinar and ductal cells (ductal log enrich=3.43, 95% CI=1.07-4.71, acinar log enrich=2.74, 95% CI=0.66-4.02) (**Figure 3b**). We further enumerated the contribution of these cell types to T1D risk by determining the cumulative posterior probability (cPPA) of fine-mapped variants overlapping cell type-specific cCREs after removing variants overlapping a more probable cell type (**see Methods**). Among broad annotation categories, distal cCREs harbored the most cumulative risk (cPPA=24.3, N_vars_=291), followed by coding exons (cPPA=7.98, N=34) and promoters (cPPA=6.63, N=55) (**Figure 3c**). When breaking down distal cCREs by cell type categories, CD4+ T cells had the most cumulative risk (cPPA=9.7, N=112), followed by exocrine cells (acinar and ductal; cPPA=6.2, N=51), monocytes (cPPA=3.1, N=54), and then endocrine cells (cPPA=2.3, N=33) (**Figure 3c**).

Given insight into cell types contributing to T1D risk, we next annotated individual T1D signals in cCREs for these cell types. Over 75% (109/141) of T1D signals contained at least one fine-mapped variant (with PPA>.01) overlapping a cCRE, and at 83% (90/109) of these signals the cCRE was further co-accessible with at least one gene promoter (**Supplementary Table 8**). For each T1D signal, we calculated the cPPA of fine-mapped variants overlapping cCREs for disease-enriched cell types. At 58 T1D signals a fine-mapped variant overlapped a CD4+ T cell cCRE, and signals with the highest cPPA in CD4+ T cells included the *CD2, IL2RA, PRF1* and *IKZF4* loci (**Figure 3d**). We also identified T1D signals with high cPPA in pancreatic acinar and ductal (exocrine) cCREs and monocyte cCREs, many of which were cell type-specific (**Figure 3d**). For example, three variants at the *GP2* locus accounted for .951 of the PPA and mapped in an acinar-specific cCRE co-accessible with the promoter of *GP2*, which encodes the major membrane glycoprotein of pancreatic zymogen granules (**Figure 3e**). Similarly, rs72802342 at the *BCAR1* locus (PPA=.30) mapped in an acinar-specific cCRE co-accessible with the *CTRB1* and *CTRB2* promoters (**Figure 3f**). We observed similar predicted mechanisms in acinar cells at the *RNLS* and *COBL* loci, as well as the novel *CEL* locus, where rs541856133 (PPA=.99) mapped in a region of broad acinar-specific accessibility although not in a cCRE directly (**Supplementary Figure 7a-c**). At *CTLA4*, variant rs3087243 (PPA=.99) mapped in an acinar-specific cCRE, although the region around the variant was also broadly accessible in regulatory T cells, in line with the specialized function of *CTLA4* in regulatory T cells^54^ (**Supplementary Figure 7d**). Exocrine cCREs harboring T1D risk variants at these loci were also largely specific relative to previous studies of accessible chromatin from stimulated immune cells^55^ and cytokine-stimulated islets^56^ except for *CTLA4* which mapped in a stimulated immune site (**Supplemental Table 8**).

### Risk variant at novel T1D locus has pancreatic ductal cell-specific effects on *CFTR*

As another example of an exocrine-specific T1D signal, at the *CFTR* locus fine-mapped variant rs7795896 (PPA=0.60) mapped in a distal cCRE highly specific to pancreatic ductal cells upstream of the *CFTR* gene (**Figure 4a**). Furthermore, the cCRE harboring rs7795896 had ductal cell-specific co-accessibility with the *CFTR* promoter in addition to several other genes (**Figure 4a**). Recessive mutations in *CFTR* cause cystic fibrosis (CF) which is often comorbid with exocrine pancreas insufficiency and CF-related diabetes (CFRD)^57^. Furthermore, carriers of *CFTR* mutations often develop chronic pancreatitis^58^. As *CFTR* has not been previously implicated in T1D, we sought to validate the mechanism of this locus. First, we determined whether rs7795896 had allele-specific activity using luciferase reporter and gel shift assays in Capan-1 cells, an established model of ductal cell function^59^. We observed both significantly reduced enhancer activity (P=3.35×10^−2^, **Figure 4b**) and reduced protein binding for the T1D risk allele (**Figure 4c**). The variant mapped in a predicted sequence motif for the ductal cell-specific transcription factor HNF1B (**Supplemental Table 6**) and overlapped a HNF1B ChIP-seq site previously identified in ductal cell models (**Supplemental Figure 8**).

To determine whether the enhancer harboring rs7795896 regulated the expression of *CFTR* in ductal cells, we used CRISPR interference (CRISPRi) to repress the activity of the enhancer (*CFTR*^Enh^) in Capan-1 cells using two independent guide RNAs. As positive and negative controls, we inactivated the *CFTR* promoter (*CFTR*^Prom^) and used a non-targeting guide RNA, respectively. RNA-seq analysis revealed a significant reduction in *CFTR* expression after enhancer inactivation (*CFTR*^Enh^ log_2_(FC)=-0.40, P=2.41×10^−3^), whereas expression of other genes co-accessible with the enhancer was unchanged (**Figure 4d**), identifying *CFTR* as a target gene of this enhancer. We next determined whether risk variants affected *CFTR* expression directly using pancreas eQTL data from GTEx^60^. Out of 13 genes tested by GTEx for association with these variants, only *CFTR* had evidence for an eQTL (P=4.31×10^−4^), and this eQTL was statistically colocalized with the T1D signal (PP_shared_=91.4%) (**Figure 4e**). The T1D risk allele C was also associated with decreased *CFTR* expression, consistent with effects on enhancer activity and TF binding. To evaluate whether the *CFTR* eQTL signal in whole pancreas tissue was driven by ductal cells, we used MuSiC^61^ to estimate cell type proportions in each GTEx pancreas RNA-seq sample (**Figure 4f, Supplemental Figure 9**). We then re-calculated eQTL association including estimated cell type proportion for each sample as an interaction term in the model, and only ductal cells had significant association (P=2.37×10^−4^) (**Figure 4g**).

As *CFTR* has been implicated in risk of pancreatic cancer^62^ and pancreatitis^63^, we finally asked whether rs7795896 was significantly associated with these phenotypes in the UK biobank^64^, FinnGen, and other GWAS^28–31^. The T1D risk allele (C) was associated with increased risk of pancreatitis (chronic pancreatitis OR=1.15, P=3.18×10^−3^; acute pancreatitis OR=1.07, P=1.15×10^−2^), pancreatic cancer (OR=1.10, P=7.85×10^−2^), and other pancreatic diseases which includes pancreatitis and pancreatic cysts (OR=1.13, P=4.72×10^−5^) (**Figure 4h**). In contrast, rs7795896 did not show evidence for association with other autoimmune diseases (all P>.05), supporting that it likely does not affect intrinsic immune cell function. Together our findings support a model in which non-coding variants regulating the activity of genes such as *CFTR* in the exocrine pancreas contribute to risk of T1D as well as pancreatic disease (**Figure 4i**).

## DISCUSSION

Population-based association studies of complex disease are a powerful tool for genetic discovery and, when coupled with cell type-resolved epigenome maps, can help reveal the cellular origins of disease. Our results represent the largest genome-wide study of T1D genetics to date, more than doubling the set of known risk signals, and provide a comprehensive resource for interrogating T1D risk mechanisms. Integration of these data with cell type-specific accessible chromatin maps both confirmed the prominent role of CD4+ T cells and implicated additional cell types in disease risk notably pancreatic acinar and ductal cells. T1D risk variants mapped to genes with specialized function in acinar and ductal cells such as *CFTR, GP2* and *CEL*, none of which have been previously implicated in T1D. Observational studies have reported exocrine pancreas abnormalities in T1D at disease onset^65^ as well as in autoantibody positive individuals^66^ and first-degree relatives of T1D^67^, but it was unknown whether this was contributing causally to disease^68,69^. Studies in zebrafish, mice and humans have demonstrated that reduced *CFTR* leads to CFRD via intra-islet inflammation and immune infiltration rather than intrinsic defects of beta cell function, and immune infiltration in the exocrine pancreas has been suggested to contribute to T1D pathogenesis^70–72^. We therefore hypothesize a causal role for gene regulation in exocrine cells in T1D, potentially mediated through immune infiltration and inflammation, which may provide novel avenues for therapeutic discovery in T1D.

## METHODS

### Genotype quality control and imputation

We compiled individual-level genotype data and summary statistics of 18,803 T1D cases and 470,876 controls of European ancestry from public sources (**Supplementary Table 1**), where T1D case cohorts were matched to population control cohorts based on genotyping array (Affymetrix, Illumina Infinium, Illumina Omni, and Immunochip) and country of origin where possible (US, British, and Ireland). For the GENIE-UK cohort, because we were unable to find a matched country of origin control cohort, we used individuals of British ancestry (defined by individuals within 1.5 interquartile range of CEU/GBR subpopulations on the first 4 PCs from PCA with European 1000 Genomes Project samples) from the University of Michigan Health and Retirement study (HRS). For non-UK Biobank cohorts, we first applied individual and variant exclusion lists (where available) to remove low quality, duplicate, or non-European ancestry samples and failed genotype calls for each cohort. For control cohorts, we also used phenotype files (where available) to remove individuals with type 2 diabetes or autoimmune diseases.

We then applied a uniform processing pipeline and used PLINK^73^ to remove variants based on (i) low frequency (MAF<1%), (ii) missing genotypes (missing>5%), (iii) violation of Hardy-Weinberg equilibrium (HWE *p*<1×10^−5^ in control cohorts and HWE *p*<1×10^−10^ in case cohorts), (iv) substantial differences in allele frequency compared to the Haplotype Reference Consortium r1.1 reference panel^74^, and (v) allele ambiguity (AT/GC variants with MAF>40%). We further removed individuals based on (i) missing genotypes (missing>5%), (ii) sex mismatch with phenotype records (het_chrX_>.2 for females and het_chrX_<.8 for males), (iii) cryptic relatedness through identity-by-descent (IBD>.2), and (iv) non-European ancestry through PCA with 1000 Genomes Project^75^ (>3 interquartile range from 25^th^ and 75^th^ percentiles of European 1KGP samples on the first 4 PCs) (**Supplementary Figure 1**). For the affected sib-pair (ASP) cohort genotyped on the Immunochip, we retained only one T1D sample from each family selected at random. For the GRID case and 1958 Birth control cohorts genotyped on the Immunochip, a portion of the cases overlapped the T1DGC or 1958 Birth cohorts genotyped on a genome-wide array. We thus used sample IDs from the phenotype files to remove these samples from the GRID and 1958 Birth cohorts and verified that no samples were duplicated between the Immunochip and genome-wide array datasets by checking IBD values. We combined data for matched case and control cohorts based on genotyping array and country of origin for imputation. We used the TOPMed Imputation Server^76,77^ to impute genotypes into the TOPMed r2 panel^7^ and removed variants based on low imputation quality (R^2^<.3). Following imputation, we implemented post-imputation filters to remove variants based on potential genotyping or imputation artifacts based on empirical R^2^ (genotyped variants with empirical R^2^<.5 and all imputed variants in at least low LD (r^2^>.3) with them).

For the UK Biobank cohort, we downloaded imputed genotype data from the UK Biobank v3 release which were imputed using a combination of the HRC and UK10K + 1000 Genomes reference panels. We used phenotype data to remove individuals of non-European descent. We then used a combination of ICD10 codes to define 1,458 T1D cases (T1D diagnosis and insulin treatment within a year of diagnosis, no T2D diagnosis). We defined controls as 362,257 individuals without diabetes (no T1D, T2D, or gestational diabetes diagnosis) or other autoimmune diseases (systemic lupus erythematosus, rheumatoid arthritis, juvenile arthritis, Sjögren syndrome, alopecia areata, multiple sclerosis, autoimmune thyroiditis, vitiligo, celiac disease, primary biliary cirrhosis, psoriasis, or ulcerative colitis). We removed variants with low imputation quality (R^2^<.3).

For the FinnGen cohort, we downloaded GWAS summary statistics for type 1 diabetes (E4_DM1_STRICT) from FinnGen freeze 2. This phenotype definition excluded individuals with type 2 diabetes from both cases and controls.

### Association testing, meta-analysis, and detection of conditional signals

We tested low-frequency and common variants (MAF>.001%) for association to T1D with firth bias reduced logistic regression using EPACTS (https://genome.sph.umich.edu/wiki/EPACTS) for non-UK Biobank cohorts or SAIGE^64^ for the UK Biobank, using genotype dosages adjusted for sex and the first four ancestry PCs. We then combined association results across matched cohorts through inverse-variance weighted meta-analysis. We used the liftOver utility to convert GRCh38/hg38 into GRCh37/hg19 coordinates for all cohorts except for the UK biobank. We removed variants that were unable to be converted, were duplicated after coordinate conversion, or were located on different chromosomes after conversion. In total, our association data contained summary statistics for 59,244,856 variants. To evaluate the extent to which genomic inflation was driven by the polygenic nature of T1D or population stratification, we used LD score regression to compare the LDSC intercept to lambda genomic control (GC). We observed an intercept of 1.08 (SE=.03) compared to a lambda GC of 1.21, suggesting that the majority of the observed inflation was driven by polygenicity rather than population stratification.

We used a threshold of P<5×10^−8^ to define genome-wide significance for primary signals, and we defined novel loci as those statistically independent (r^2^<.01) from reported index variants from previous T1D association studies. For all cohorts except for FinnGen, we performed exact conditional analyses on lead index variants to identify conditionally independent signals and used a locus-wide threshold of P<1×10^−5^ to define significance. For genomic regions with multiple known signals within close proximity, we conditioned on index variants from both signals. We iterated through this process for each locus until there were no remaining significant signals at the locus-wide threshold.

### Fine mapping of distinct association signals

We constructed LD-based genetic credible sets of variants for 141 signals at 89 known and novel loci excluding the MHC locus for complex LD structure and *ICOSLG*, for which we were unable to find imputed proxy variants in our dataset. For the main signals at known loci, we defined credible set variants by taking all variants in at least low LD (r^2^>.1) with newly identified index variants within a 5 Mb window. For both novel and conditional signals, we used the most significant variant at the signal and the same credible set definition. We used effect size and standard error estimates to calculate approximate Bayes factors^9^ (ABF) for each variant; at signals with multiple distinct association signals, we derived values from the corresponding conditional analysis. We then calculated the posterior probability of association (PPA) for each variant by dividing its ABF by the sum of ABF for all variants in the signal’s credible set. To derive 99% credible sets for each signal, we sorted variants for each signal by descending PPA and retained variants that added up to a cumulative PPA>0.99. To verify that variant coverage across different imputation panels did not affect fine mapping, we calculated the effective sample size for all credible set variants. There were only 9 credible set variants in total with <50% of the maximum effective sample size, all of which had PPA<.01, and we did not further filter these variants.

### GWAS correlation analyses

We used LD score regression (version 1.0.1) to estimate genome-wide genetic correlations between T1D and immune diseases^25–31,41,42^, other diseases^32–40,64,78,79^, and non-disease traits^43–50,80–88^, using European subsets of GWAS where applicable. For acute pancreatitis, chronic pancreatitis, and pancreatic cancer, we used inverse variance weighted meta-analysis to combine SAIGE analysis results from the UK biobank^64^ (PheCodes 577.1, 577.2, and 157) and FinnGen (K11_ACUTPANC, K11_CHRONPANC, C3_PANCREAS_EXALLC). We used pre-computed European 1000 Genomes LD scores to calculate correlation estimates (r_g_) and standard errors. We then corrected p-values for multiple tests using FDR correction, considering traits with FDR<.1 as significant. We also performed genetic correlation analyses using a version of the T1D meta-analysis excluding the Immunochip cohorts and observed highly similar results.

### Generation of snATAC-seq libraries

#### Combinatorial indexing single cell ATAC-seq (snATAC-seq/sci-ATAC-seq)

snATAC-seq was performed as described previously^89,90^ with several modifications as described below. For the islet samples, approximately 3,000 islet equivalents (IEQ, roughly 1,000 cells each) were resuspended in 1 mL nuclei permeabilization buffer (10mM Tris-HCL (pH 7.5), 10mM NaCl, 3mM MgCl_2_, 0.1% Tween-20 (Sigma), 0.1% IGEPAL-CA630 (Sigma) and 0.01% Digitonin (Promega) in water) and homogenized using 1mL glass dounce homogenizer with a tight-fitting pestle for 15 strokes. Homogenized islets were incubated for 10 min at 4°C and filtered with 30 µm filter (CellTrics). For the pancreas samples, frozen tissue was pulverized with a mortar and pestle while frozen and immersed in liquid nitrogen. Approximately 22 mg of pulverized tissue was then transferred to an Eppendorf tube and resuspended in 1 mL of cold permeabilization buffer for 10 minutes on a rotator at 4°C. Permeabilized sample was filtered with a 30µm filter (CellTrics), and the filter was washed with 300 µL of permeabilization buffer to increase nuclei recovery.

Once permeabilized and filtered, nuclei were pelleted with a swinging bucket centrifuge (500 x g, 5 min, 4°C; 5920R, Eppendorf) and resuspended in 500 µL high salt tagmentation buffer (36.3 mM Tris-acetate (pH = 7.8), 72.6 mM potassium-acetate, 11 mM Mg-acetate, 17.6% DMF) and counted using a hemocytometer. Concentration was adjusted to 4500 nuclei/9 µl, and 4,500 nuclei were dispensed into each well of a 96-well plate. Glycerol was added to the leftover nuclei suspension for a final concentration of 25 % and nuclei were stored at −80°C. For tagmentation, 1 µL barcoded Tn5 transposomes^90^ were added using a BenchSmart™ 96 (Mettler Toledo), mixed five times and incubated for 60 min at 37°C with shaking (500 rpm). To inhibit the Tn5 reaction, 10 µL of 40 mM EDTA were added to each well with a BenchSmart™ 96 (Mettler Toledo) and the plate was incubated at 37°C for 15 min with shaking (500 rpm). Next, 20 µL 2 x sort buffer (2 % BSA, 2 mM EDTA in PBS) were added using a BenchSmart™ 96 (Mettler Toledo). All wells were combined into a FACS tube and stained with 3 µM Draq7 (Cell Signaling). Using a SH800 (Sony), 20 nuclei were sorted per well into eight 96-well plates (total of 768 wells) containing 10.5 µL EB (25 pmol primer i7, 25 pmol primer i5, 200 ng BSA (Sigma))^90^. Preparation of sort plates and all downstream pipetting steps were performed on a Biomek i7 Automated Workstation (Beckman Coulter). After addition of 1 µL 0.2% SDS, samples were incubated at 55 °C for 7 min with shaking (500 rpm). We added 1 µL 12.5% Triton-X to each well to quench the SDS and 12.5 µL NEBNext High-Fidelity 2× PCR Master Mix (NEB). Samples were PCR-amplified (72 °C 5 min, 98 °C 30 s, (98 °C 10 s, 63 °C 30 s, 72 °C 60 s) × 12 cycles, held at 12 °C). After PCR, all wells were combined. Libraries were purified according to the MinElute PCR Purification Kit manual (Qiagen) using a vacuum manifold (QIAvac 24 plus, Qiagen) and size selection was performed with SPRI Beads (Beckmann Coulter, 0.55x and 1.5x). Libraries were purified one more time with SPRI Beads (Beckmann Coulter, 1.5x). Libraries were quantified using a Qubit fluorimeter (Life technologies) and the nucleosomal pattern was verified using a TapeStation (High Sensitivity D1000, Agilent). The library was sequenced on a HiSeq2500 sequencer (Illumina) using custom sequencing primers, 25% spike-in library and following read lengths: 50 + 43 + 40 + 50 (Read1 + Index1 + Index2 + Read2).

#### Droplet-based 10X single cell ATAC-seq (scATAC-seq)

10X scATAC-seq protocol from 10x Genomics was followed: Chromium SingleCell ATAC ReagentKits UserGuide (CG000209, Rev A). Cryopreserved PBMC samples were thawed in 37°C water bath for 2 min and followed ‘PBMC thawing protocol’ in the UserGuide. After thawing cells, the pellets were resuspended again in 1 mL chilled PBS (with 0.04% PBS) and filtered with 50 μm CellTrics (04-0042-2317, Sysmex). The cells were centrifuged (300g, 5 min, 4°C) and permeabilized with 100 μl of chilled lysis buffer (10 mM Tris-HCl pH 7.4, 10 mM NaCl, 3 mM MgCl2, 0.1% Tween-20, 0.1% IGEPAL-CA630, 0.01% digitonin and 1% BSA). The samples were incubated on ice for 3 min and resuspended with 1mL chilled wash buffer (10 mM Tris-HCl pH 7.4, 10 mM NaCl, 3 mM MgCl2, 0.1% Tween-20 and 1% BSA). After centrifugation (500g, 5 min, 4°C), the pellets were resuspended in 100 µL of chilled Nuclei buffer (2000153, 10x Genomics). The nuclei concentration was adjusted between 3,000 to 7,000 per μl and 15,300 nuclei which targets 10,000 nuclei was used for the experiment. For pancreas tissue (pulverized as described above), approximately 31.7 mg of pulverized tissue was transferred to a LoBind tube (Eppendorf) and resuspended in 1 mL of cold permeabilization buffer (10mM Tris-HCL (pH 7.5), 10mM NaCl, 3mM MgCl_2_, 0.1% Tween-20 (Sigma), 0.1% IGEPAL-CA630 (Sigma), 0.01% Digitonin (Promega) and 1% BSA (Proliant 7500804) in water) for 10 min on a rotator at 4°C. Permeabilized nuclei were filtered with 30 µm filter (CellTrics). Filtered nuclei were pelleted with a swinging bucket centrifuge (500 x g, 5 min, 4°C; 5920R, Eppendorf) and resuspended in 1 mL Wash buffer (10mM Tris-HCL (pH 7.5), 10mM NaCl, 3mM MgCl_2_, 0.1% Tween-20, and 1% BSA (Proliant 7500804) in molecular biology-grade water). Nuclei wash was repeated once. Next, washed nuclei were resuspended in 30 µL of 1X Nuclei Buffer (10X Genomics). Nuclei were counted using a hemocytometer, and finally the nuclei concentration was adjusted to 3,000 nuclei/µl. 15,360 nuclei were used as input for tagmentation.

Nuclei were diluted to 5 μl with 1X Nuclei buffer (10x Genomics) and, mixed with ATAC buffer (10x Genomics) and ATAC enzyme (10x Genomics) for tagmentation (60 min, 37°C). Single cell ATAC-seq libraries were generated using the (Chromium Chip E Single Cell ATAC kit (10x Genomics, 1000086) and indexes (Chromium i7 Multiplex Kit N, Set A, 10x Genomics, 1000084) following manufacturer instructions. Final libraries were quantified using a Qubit fluorimeter (Life technologies) and the nucleosomal pattern was verified using a TapeStation (High Sensitivity D1000, Agilent). Libraries were sequenced on a NextSeq 500 and HiSeq4000 sequencer (Illumina) with following read lengths: 50 + 8 + 16 + 50 (Read1 + Index1 + Index2 + Read2).

### Single cell chromatin accessibility data processing

Prior to read alignment, we used trim_galore (version 0.4.4) to remove adapter sequences from reads using default parameters. We aligned reads to the hg19 reference genome using bwa mem^91^ (version 0.7.17; parameters: ‘-M −C’) and removed low mapping quality (MAPQ<30), secondary, unmapped, and mitochondrial reads using samtools^92^. To remove duplicate sequences on a per-barcode level, we used the MarkDuplicates tool from picard (parameters: ‘BARCODE_TAG’). For each tissue and snATAC-seq technology, we used log-transformed read depth distributions from each experiment to determine a threshold separating real cell barcodes from background noise. We used 500 total reads (passing all filters) as the cutoff for combinatorial barcoding snATAC and between 2,300 and 4,000 total reads, as well as at least 0.3 fraction of reads in peaks for 10x snATAC-seq experiments (**Supplemental Figure 5a**).

### Single cell chromatin accessibility clustering

We identified snATAC-seq clusters using a previously described pipeline with a few modifications. For each experiment, we first constructed a counts matrix consisting of read counts in 5 kb windows for each cell. Using scanpy^93^, we normalized cells to a uniform read depth and log-transformed counts. We extracted highly variable (*hv*) windows (parameters: ‘min_mean=.01, min_disp=.25’) and regressed out the total log-transformed read depth within *hv* windows (usable counts). We then merged datasets from the same tissue and performed PCA to extract the top 50 PCs. We used Harmony^94^ to correct the PCs for batch effects across experiments, using categorical covariates such as donor-of-origin (all tissues), biological sex (PBMCs), and snATAC-seq assay technology (pancreas). We used the corrected components to construct a 30 nearest neighbor graph using the cosine metric, which we used for UMAP dimensionality reduction (parameters: ‘min_dist=.3’) and clustering with the Leiden algorithm^95^ (parameters: ‘resolution=1.5’).

Prior to combining cells across all tissues, we performed iterative clustering to identify and remove cells with aberrant quality metrics. First, we identified and remove clusters of cells with lower quality metrics (islets: 948, pancreas: 2,588, PBMCs: 5,268 cells removed total), including lower usable counts or fraction of reads in peaks. Next, after removing the low-quality cells and repeating the previous clustering steps, we sub-clustered the resulting main clusters at high resolution (parameters: ‘resolution=3.0’) to identify sub-clusters containing potential doublets (islets: 886, pancreas: 4,495, PBMCs: 5,844 cells removed total). We noted that these sub-clusters tended to have higher average usable counts, promoter usage, and accessibility at more than one marker gene promoter. After removing 20,029 low-quality or potential doublet cells, we performed one final round of clustering using experiments from all tissues, including tissue-of-origin as another covariate. We further removed 672 cells mapping to improbable cluster assignments (islet or pancreatic cells in PBMC clusters or vice versa). After all filters, we ended up with 131,554 cells mapping to 28 distinct clusters with consistent representation across samples from the same tissue (**Supplemental Figure 5b**). We cataloged known marker genes for each cell type and assessed gene accessibility (sum of read counts across each gene body) to assign labels to each cluster.

### Single cell chromatin accessibility analyses

We identified chromatin accessibility peaks with MACS2^96^ by calling peaks on aggregated reads from each cluster. In brief, we extracted reads from all cells within a given cluster, shifted reads aligned to the positive strand by +4 bp and reads aligned to the negative strand by −5 bp, and centered the reads. We then used MACS2 to call peaks (parameters: ‘--nomodel −-keep-dup-all’) and removed peaks overlapping ENCODE blacklisted regions^97^. We then merged peaks from all 28 clusters with bedtools^98^ to create a consistent set of 448,142 regulatory elements for subsequent analyses.

To compare accessible chromatin profiles from snATAC-seq to those from bulk ATAC-seq on FACS purified cell types, we reprocessed published ATAC-seq data from sorted pancreatic^99^ and unstimulated immune cells^55^. We created pseudobulk profiles from the snATAC-seq data for each donor and cluster, retaining those that contained information from at least 50 cells. We then extracted read counts in the 448,142 merged peaks for all sorted and pseudobulk profiles. We used PCA to extract the top 20 principal components and used UMAP for dimensionality reduction and visualization (parameters: ‘min_dist=.5, n_neighbors=80’).

To identify cluster-specific peaks, we used logistic regression models for each peak treating each cell as an individual data point. For each model, we used cluster assignment and covariates such as donor-of-origin and the log usable count as predictors and binary accessibility of the peak as the outcome to calculate t-statistics (t-stats) for specificity. For a given cluster, we defined cluster-specific peaks by taking the top 1000 peaks with the highest t-stats, after first filtering out peaks which also had high t-stats for other clusters (peak t-stat>90^th^ percentile of all t-stats for the given cluster in more than 2 other clusters). We then used GREAT^12^ to annotate peaks and summarize linked genes in the form of gene ontology terms for the set of cluster-specific peaks as compared to all merged peaks.

We estimated TF motif enrichment z-scores for each cell using chromVAR^13^ (version 1.5.0) by following the steps outlined in the user manual. First, we constructed a sparse binary matrix encoding read overlap with merged peaks for each cell. For each merged peak, we estimated the GC content bias based on the hg19 human reference genome to obtain a set of matched background peaks. To ensure a motif enrichment value for each cell, we did not apply any additional filters based on total reads or the fraction of reads in peaks. Next, using 580 TF motifs within the JASPAR 2018 CORE vertebrate (non-redundant) set^14^, we computed GC bias-corrected enrichment z-scores (chromVAR deviation scores) for each cell. To extract highly variable TF motifs, we computed the enrichment variability of each motif across all cells and used the median as the cutoff. For each cluster, we then computed the average TF motif enrichment z-score across all cells in the cluster.

We used Cicero^24^ (version 1.3.3) to calculate co-accessibility scores between pairs of peaks for each cluster. As in the single cell motif enrichment analysis, we started from a sparse binary matrix. For each cluster, we only retained merged peaks that overlapped peaks from the cluster. Within each cluster, we aggregated cells based on the 50 nearest neighbors and used cicero to calculate co-accessibility scores, using a 1 Mb window size and a distance constraint of 500 kb. We then defined promoters as ±500 bp from the TSS of protein coding transcripts to annotate co-accessibility links between distal and promoter peaks.

### GWAS enrichment analyses

We used LD score regression^100^ to calculate genome-wide enrichment z-scores for 32 diseases and traits including T1D. We obtained GWAS summary statistics for autoimmune and inflammatory diseases (immune-related)^25–31,41,42^, other diseases^32–40^, and quantitative endophenotypes^43–52^, and where necessary, we filled in variant IDs and alleles. Using the ‘munge_sumstats.py’ script, we converted summary statistics to the standard format for LD score regression. For each cluster, we used overlap with chromatin accessibility peaks as a binary annotation for variants. We also created a background annotation using merged peaks across all clusters. Then, we computed annotation-specific LD scores by following the instructions for creating partitioned LD scores. We used stratified LD score regression^53^ to estimate enrichment coefficient z-scores for each annotation relative to the background, which we defined as merged peaks across all clusters combined with the annotations in the baseline-LD model (version 2.2). Based on the enrichment z-scores, we computed one-sided p-values to assess significance and corrected for multiple tests using the Benjamini-Hochberg procedure^101^. We also calculated GWAS enrichment z-scores for T1D using a version of the meta-analysis excluding the Immunochip cohorts and observed highly similar enrichment results. We used fgwas to estimate enrichment within cell type-specific cCREs using 2000 variants per window.

### Annotating cell type mechanisms of variants at fine mapped signals

We first annotated fine mapped variants with PPA>1% using broad genomic annotations. We defined “coding” as coding exons of protein coding genes, “promoter” as ±500 bp from the TSS of protein coding transcripts, and “distal” as peaks in any cell type that did not overlap promoter regions. We then assigned variants to each group without replacement, in the priority coding>promoter>distal. To then further breakdown distal variants, we assigned clusters to cell type groups (CD4 T cell: naïve CD4 T, activated CD4 T, regulatory T; CD8 T cell: naïve CD8 T, activated CD8 T, pancreatic CD8 T; NK cell: adaptive and cytotoxic NK; B cell: naïve and memory B; monocyte/ MΦ: classical and non-classical monocyte, pancreatic macrophage; dendritic: conventional and plasmacytoid dendritic; other cell: megakaryocyte, endothelial, activated and quiescent stellate; exocrine: acinar and ductal; endocrine: alpha, beta, delta, and gamma) and created merged peak annotations for each group. We then assigned variants to each cell type group without replacement, prioritizing groups in order based on their cumulative PPA.

### Luciferase reporter assay

To test for allelic differences in enhancer activity at rs7795896, we cloned human DNA sequences (Coriell) containing the reference or alternate allele upstream of the minimal promoter in the luciferase reporter vector pGL4.23 (Promega) in the forward direction using the restriction enzymes SacI and KpnI. We then created a construct containing the alternate allele using the NEB Q5 SDM kit (New England Biolabs). The primer sequences used were:

Cloning FWD_P1 TAGCGGTACCTAATGGGAAATCATGCCAACC Cloning FWD_P2 AATAGAGCTCATGTGTGTGTGCTGGGATGT

We grew Capan-1 cells (ATCC) to approximately 70% confluency in 6-well dishes according to ATCC culture recommendations. We co-transfected cells with either the experimental or empty vector and pRL-SV40. We then lysed cells 48 hours post transfection and assayed them using the Dual-Luciferase Reporter System (Promega). We normalized Firefly activity to Renilla activity and expressed normalized results as fold change compared to the luciferase activity of the empty vector. We used a two-sided t-test to compare the luciferase activity between the two alleles.

### Electrophoretic mobility shift assay

We ordered 5’ biotinylated and unlabeled (cold) oligos with the reference and alternate alleles from Integrated DNA Technologies. We annealed oligos with an equivalent volume of equimolar complementary oligo in a binding buffer containing 10mM Tris pH 8.0, 50mM NaCl, and 1mM EDTA at 95°C for 5 minutes and cooled them gradually to room temperature before further use.

C oligo: (5’ biotin)CAATTAGATGTAACTCATTAACATTAGAAAAA T oligo: (5’ biotin)CAATTAGATGTAACTTATTAACATTAGAAAAA

We carried out binding reactions using the LightShift Chemiluminescent EMSA kit (Thermo Fisher) according to manufacturer’s instructions with the following adjustments: 100 fmol of biotinylated probe per reaction and 20 pmol of non-biotinylated “cold” probe in competition reactions. We used approximately 16 ug of nuclear protein extract from Capan-1 cells purified using NE-PER Nuclear and Cytoplasmic Extraction Reagents (Thermo Fisher) per binding reaction.

### CRISPR inactivation of enhancer element

We maintained HEK293T cells in DMEM containing 100 units/mL penicillin and 100 mg/mL streptomycin sulfate supplemented with 10% fetal bovine serum (FBS). To generate CRISPRi expression vectors, we designed guide RNA sequences to target the enhancer containing rs7795896 or the *CFTR* promoter. These guides, as well as a non-targeting control, were placed downstream of the human U6 promoter in the pLV hU6-sgRNA hUbC-dCas9-KRAB-T2a-Puro backbone (Addgene, #71236). The guide RNA sequences were:

rs7795896 enhancer guide 1 GTAGTTGGCTTCCTCAGTAAG

rs7795896 enhancer guide 2 GAACAGTATGATTTACGTAA

*CFTR* promoter GCGCCCGAGAGACCATGCAG

Non-targeting control GTGACGTGCACCGCGGTGTG

We generated high-titer lentiviral supernatants by co-transfection of the resulting plasmid and lentiviral packaging constructs into HEK293T cells. Specifically, we co-transfected CRISPRi vectors with the pCMV-R8.74 (Addgene, #22036) and pMD2.G (Addgene, #12259) expression plasmids into HEK293T cells using a 1mg/mL PEI solution (Polysciences). We collected lentiviral supernatants at 48 hours and 72 hours after transfection and concentrated lentiviruses by ultracentrifugation for 120 minutes at 19,500 rpm using a Beckman SW28 ultracentrifuge rotor at 4°C.

We obtained Capan-1 pancreatic ductal adenocarcinoma cell lines from ATCC and cultured them using Iscove’s Modified Dulbecco’s Media with 20% fetal bovine serum, 100 units/mL penicillin, and 100 mg/mL streptomycin sulfate. 24 hours prior to infection, we passaged cells into a 6-well plate at a density of 650,000 cells per well. The following day, we added fresh media containing 5ug/mL polybrene and 5uL/mL concentrated CRISPRi lentivirus to each well. We incubated the cells at 37°C for 30 minutes and then spun them in a centrifuge for 1 hour at 30°C at 950 × g. 6 hours later, we replaced viral media with fresh base culture media and left the cells to recover. After 48 hours, we replaced media daily with the addition of 2ug/mL puromycin for a further 72 hours. We then harvested infected cells and isolated RNA using the RNeasy® Micro Kit (Qiagen) according to the manufacturer instructions.

### Differential analysis of CRISPR inactivation experiments

We used STAR (version 2.7.3a) to map reads to the hg19 genome using ENCODE standard options (parameters: ‘--outFilterType BySJout --outFilterMultimapNmax 20 --alignSJoverhangMin 8 --alignSJDBoverhangMin 1 --outFilterMismatchNmax 999 –outFilterMismatchNoverReadLmax 0.04 --alignIntronMin 20 --alignIntronMax 1000000 --alignMatesGapMax 1000000’). We then used featureCounts (version 1.6.4) to count the number of uniquely mapped reads mapping to genes in GENCODE v19 (parameters: ‘-Q 30 -p -B -s 2 --ignoreDup’). We used DESeq2 to evaluate differential mRNA expression between either the *CFTR* enhancer (pooled data from both guides), or promoter inactivation versus the non-targeting guide.

### Colocalization and deconvolution of the pancreas *CFTR* eQTL

We obtained GTEx consortium release v7^60^ eQTL summary statistics for pancreas tissue from 220 samples and used effect size and standard error estimates to calculate Bayes factors^9^ for each variant. Where a T1D-associated variant had evidence for a pancreas eQTL, we considered all variants in a 500kb window around the T1D GWAS index variant, and used the coloc^102^ package to calculate the probability that the variants driving T1D association and eQTL signals were shared. We considered signals as colocalized based on the probability that they were shared (PP_shared_>.9).

We downloaded and re-processed a published pancreas single cell RNA-seq dataset^103^ of 12 islet donors. After re-processing and generating a counts matrix with the 10x Genomics cellranger (version 3.0.0) pipeline, we first used scanpy^93^ and filtered out 1) cells with <500 genes expressed, 2) cells with >20% mitochondrial reads, or 3) genes expressed in <3 cells. To ensure clustering would not be affected by read depth, we normalized the total counts per cell to 10k and subsequently log-normalized the resulting counts. We identified highly variable genes (hvgs) based on mean expression and dispersion with (parameters: ‘min_mean=.005, max_mean=6, min_disp=.1’). We then extracted counts for hvgs and regressed out the total read count within the hvgs. After dimensionality reduction with PCA, we used harmony^94^ with default parameters to correct for batch effects due to donor. We used the top 30 corrected PCs for graph-based clustering with the leiden algorithm^95^ (parameters: ‘resolution=1.25’) and visualization on reduced dimensions with UMAP^104^ (parameters: ‘min_dist=.3’). To assign cell types to each cluster, we used well-established marker genes from literature and labelled 18,279 cells.

We used MuSiC^61^ to estimate the proportions of major pancreatic cell types (acinar, duct, stellate, alpha, beta, delta, gamma) in each pancreas sample from the GTEx v7 release. As input, we used raw count matrices of the islet scRNA-seq and GTEx v7 pancreas samples and cell type labels from the analysis of the former dataset. For each cell type, we used the proportion as an interaction term and constructed linear models of CFTR expression (TMM normalized) as a function of the interaction between genotype dosage and cell type proportion, accounting for covariates used by GTEx including sex, sequencing platform, 3 genotype PCs, and 28 inferred PCs from the expression data. From the original 30 inferred PCs, we excluded inferred PCs 2 and 3 because they were highly correlated (Spearman’s ρ>.7) with acinar cell proportion.

### Phenotype associations at *CFTR* variant

We tested for association of the T1D index variant rs7795896 at *CFTR* to pancreatic and autoimmune disease phenotypes. For acute pancreatitis, chronic pancreatitis, and pancreatic cancer, we used inverse variance weighted meta-analysis to combine SAIGE analysis results from the UK biobank^64^ (PheCodes 577.1, 577.2, and 157) and FinnGen (K11_ACUTPANC, K11_CHRONPANC, C3_PANCREAS_EXALLC). As mutations that cause cystic fibrosis (CF) map to this locus, which are risk factors for pancreatitis and pancreatic cancer, we determined the impact of the most common CF mutation F508del/rs199826652 on the association results for rs7795896. For T1D, we tested for association of rs7795896 conditional on F508del/rs199826652 in all cohorts except for FinnGen and observed no evidence for a difference in T1D association. For pancreatitis and pancreatic cancer, we identified F508del/rs199826652 carriers in UK Biobank and repeated the association analysis for these phenotypes in UK biobank data after removing these individuals and observed no evidence of a change in the effect of rs7795896.

## CODE AVAILABILITY

Code used for processing snATAC-seq datasets and clustering cells is available at https://github.com/kjgaulton/pipelines/tree/master/T1D_snATAC_pipeline.

## DATA AVAILABILITY

Summary statistics and fine mapping credible sets for T1D GWAS will be available in the GWAS catalog and in the T1D Knowledge Portal (http://t1d.hugeamp.org). Raw data files for snATAC-seq will be deposited to GEO, and processed data files for snATAC-seq will be available through the Diabetes Epigenome Atlas (https://www.diabetesepigenome.org/).

## ACKNOWLEDGEMENTS

This work was supported by NIH grants DK112155, DK120429 and DK122607 to K.J.G and M.S., and T32 GM008666 to R.G. We thank Samantha Kuan in the Ren Lab at the LICR for assistance with sequencing.

nPOD: This research was performed with the support of the Network for Pancreatic Organ donors with Diabetes (nPOD; RRID:SCR_014641), a collaborative type 1 diabetes research project sponsored by JDRF (nPOD: 5-SRA-2018-557-Q-R) and The Leona M. & Harry B. Helmsley Charitable Trust (Grant #2018PG-T1D053). The content and views expressed are the responsibility of the authors and do not necessarily reflect the official view of nPOD. Organ Procurement Organizations (OPO) partnering with nPOD to provide research resources are listed at http://www.jdrfnpod.org/for-partners/npod-partners/.

DCCT/EDIC: The Diabetes Control and Complications Trial (DCCT) and its follow-up the Epidemiology of Diabetes Interventions and Complications (EDIC) study were conducted by the DCCT/EDIC Research Group and supported by National Institute of Health grants and contracts and by the General Clinical Research Center Program, NCRR. The data from the DCCT/EDIC study were supplied by the NIDDK Central Repositories.

GENIE: The Genetics of Nephropathy, an International Effort (GENIE) study was conducted by the GENIE Investigators and supported by the National Institute of Diabetes and Digestive and Kidney Diseases (NIDDK). The data from the GENIE study reported here were supplied by the GENIE investigators from the Broad Institute of MIT and Harvard, Queens University Belfast and the University of Dublin.

GoKinD: The Genetics of Kidneys in Diabetes (GoKinD) Study was conducted by the GoKinD Investigators and supported by the Juvenile Diabetes Research Foundation, the CDC, and the Special Statutory Funding Program for Type 1 Diabetes Research administered by the National Institute of Diabetes and Digestive and Kidney Diseases (NIDDK). The data [and samples] from the GoKinD study were supplied by the NIDDK Central Repositories. This manuscript was not prepared in collaboration with Investigators of the GoKinD study and does not necessarily reflect the opinions or views of the GoKinD study, the NIDDK Central Repositories, or the NIDDK.

T1DGC: This research utilizes resources provided by the Type 1 Diabetes Genetics Consortium (T1DGC), a collaborative clinical study sponsored by the National Institute of Diabetes and Digestive and Kidney Diseases (NIDDK), National Institute of Allergy and Infectious Diseases (NIAID), National Human Genome Research Institute (NHGRI), National Institute of Child Health and Human Development (NICHD), and the Juvenile Diabetes Research Foundation International (JDRF) and supported by U01 DK062418. The UK case series collection was additionally funded by the JDRF and Wellcome Trust and the National Institute for Health Research Cambridge Biomedical Centre, at the Cambridge Institute for Medical Research, UK (CIMR), which is in receipt of a Wellcome Trust Strategic Award (079895). The data from the T1DGC study were supplied by dbGAP. This manuscript was not prepared in collaboration with Investigators of the T1DGC study and does not necessarily reflect the opinions or views of the T1DGC study or the study sponsors.

T1DGC (ASP/UK GRID): This research was performed under the auspices of the Type 1 Diabetes Genetics Consortium, a collaborative clinical study sponsored by the National Institute of Diabetes and Digestive and Kidney Diseases (NIDDK), National Institute of Allergy and Infectious Diseases (NIAID), National Human Genome Research Institute (NHGRI), National Institute of Child Health and Human Development (NICHD), and Juvenile Diabetes Research Foundation International (JDRF).

WTCCC: This study makes use of data generated by the Wellcome Trust Case Control Consortium. A full list of the investigators who contributed to the generation of the data is available from www.wtccc.org.uk. Funding for the project was provided by the Wellcome Trust under award 076113.

UK Biobank: Data from the UK Biobank was accessed under application 24058.

FinnGen: We want to acknowledge the participants and investigators of the FinnGen study.

CSGNM: We thank the participants of the Trinity Student Study. This study was supported by the Intramural Research Programs of the National Institutes of Health, the National Human Genome Research Institute, and the Eunice Kennedy Shriver National Institute of Child Health and Development.

NIMH Schizophrenia Controls: Funding support for the Genome-Wide Association of Schizophrenia Study was provided by the National Institute of Mental Health (R01 MH67257, R01 MH59588, R01 MH59571, R01 MH59565, R01 MH59587, R01 MH60870, R01 MH59566, R01 MH59586, R01 MH61675, R01 MH60879, R01 MH81800, U01 MH46276, U01 MH46289 U01

MH46318, U01 MH79469, and U01 MH79470) and the genotyping of samples was provided through the Genetic Association Information Network (GAIN). The datasets used for the analyses described in this manuscript were obtained from the database of Genotypes and Phenotypes (dbGaP) found at http://www.ncbi.nlm.nih.gov/gap through dbGaP accession number phs000021.v3.p2. Samples and associated phenotype data for the Genome-Wide Association of Schizophrenia Study were provided by the Molecular Genetics of Schizophrenia Collaboration (PI: Pablo V. Gejman, Evanston Northwestern Healthcare (ENH) and Northwestern University, Evanston, IL, USA).

Neurodevelopmental Genomics: Support for the collection of the data for Philadelphia Neurodevelopment Cohort (PNC) was provided by grant RC2MH089983 awarded to Raquel Gur and RC2MH089924 awarded to Hakon Hakonarson. Subjects were recruited and genotyped through the Center for Applied Genomics (CAG) at The Children’s Hospital in Philadelphia (CHOP). Phenotypic data collection occurred at the CAG/CHOP and at the Brain Behavior Laboratory, University of Pennsylvania.

eMERGE Network: Group Health Cooperative/University of Washington – Funding support for Alzheimer’s Disease Patient Registry (ADPR) and Adult Changes in Thought (ACT) study was provided by a U01 from the National Institute on Aging (Eric B. Larson, PI, U01AG006781). A gift from the 3M Corporation was used to expand the ACT cohort. DNA aliquots sufficient for GWAS from ADPR Probable AD cases, who had been enrolled in Genetic Differences in Alzheimer’s Cases and Controls (Walter Kukull, PI, R01 AG007584) and obtained under that grant, were made available to eMERGE without charge. Funding support for genotyping, which was performed at Johns Hopkins University, was provided by the NIH (U01HG004438). Genome-wide association analyses were supported through a Cooperative Agreement from the National Human Genome Research Institute, U01HG004610 (Eric B. Larson, PI). Mayo Clinic – Samples and associated genotype and phenotype data used in this study were provided by the Mayo Clinic. Funding support for the Mayo Clinic was provided through a cooperative agreement with the National Human Genome Research Institute (NHGRI), Grant #: UOIHG004599; and by grant HL75794 from the National Heart Lung and Blood Institute (NHLBI). Funding support for genotyping, which was performed at The Broad Institute, was provided by the NIH (U01HG004424). Marshfield Clinic Research Foundation – Funding support for the Personalized Medicine Research Project (PMRP) was provided through a cooperative agreement (U01HG004608) with the National Human Genome Research Institute (NHGRI), with additional funding from the National Institute for General Medical Sciences (NIGMS) The samples used for PMRP analyses were obtained with funding from Marshfield Clinic, Health Resources Service Administration Office of Rural Health Policy grant number D1A RH00025, and Wisconsin Department of Commerce Technology Development Fund contract number TDF FYO10718. Funding support for genotyping, which was performed at Johns Hopkins University, was provided by the NIH (U01HG004438). Northwestern University – Samples and data used in this study were provided by the NUgene Project (www.nugene.org). Funding support for the NUgene Project was provided by the Northwestern University’s Center for Genetic Medicine, Northwestern University, and Northwestern Memorial Hospital. Assistance with phenotype harmonization was provided by the eMERGE Coordinating Center (Grant number U01HG04603). This study was funded through the NIH, NHGRI eMERGE Network (U01HG004609). Funding support for genotyping, which was performed at The Broad Institute, was provided by the NIH (U01HG004424).

Vanderbilt University -Funding support for the Vanderbilt Genome-Electronic Records (VGER) project was provided through a cooperative agreement (U01HG004603) with the National Human Genome Research Institute (NHGRI) with additional funding from the National Institute of General Medical Sciences (NIGMS). The dataset and samples used for the VGER analyses were obtained from Vanderbilt University Medical Center’s BioVU, which is supported by institutional funding and by the Vanderbilt CTSA grant UL1RR024975 from NCRR/NIH. Funding support for genotyping, which was performed at The Broad Institute, was provided by the NIH (U01HG004424). Assistance with phenotype harmonization and genotype data cleaning was provided by the eMERGE Administrative Coordinating Center (U01HG004603) and the National Center for Biotechnology Information (NCBI). The datasets used for the analyses described in this manuscript were obtained from dbGaP at http://www.ncbi.nlm.nih.gov/gap through dbGaP accession number phs000360.v3.p1.

This manuscript was not prepared in collaboration with investigators of these studies and does not necessarily reflect the opinions or views of the DCCT/EDIC, GENIE, GoKinD, T1DGC, WTCCC, studies or study groups, the NIDDK Central Repositories, the NIH, or the study sponsors.

## AUTHOR CONTRIBUTIONS

K.J.G and J.C. designed the study and wrote the manuscript. J.C. performed the genetic association and single cell accessible chromatin analyses. R.G., M.O. and S.Huang performed molecular experiments of enhancer and variant function. J.Y.H and M.M. generated single cell accessible chromatin data. P.B. and K.K. contributed to data analysis. D.U.G and S.P. supervised the generation of single cell accessible chromatin and contributed to data interpretation and analyses. M.S. supervised experiments related to enhancer function and contributed to data interpretation. S.Heller and A.K. contributed to interpretation of experimental data.

## Notes

### Competing Interest Statement

KJG does consulting for Genentech and holds stock in Vertex Pharmaceuticals

