## Supplementary Figures for "Large-scale genetic association and single cell accessible chromatin mapping defines cell type-specific mechanisms of type 1 diabetes risk"

### Figure S1

**Variant harmonization with HRC.** Update strand, alleles, and genomic coordinates of variants to reflect HRC/hg19. Remove variants with substantial difference in allele frequency compared to HRC (>20%, except for MHC region for case cohorts) or ambiguous alleles (AT/GC variants with MAF>40%).

**Variant level QC.** Remove autosomal variants with missing genotypes (>5%), low MAF (<1%), deviating from Hardy-Weinberg equilibrium ($P<1\times10^{-5}$ for control cohorts and $P<1\times10^{-10}$ for case cohorts), or in variant exclusion lists (where available).

**Sample level QC.** Remove samples with missing genotypes (>5% missing), mis-annotated sex (chrX homozygosity <.8 for males or >.2 for females), related samples (IBD >.2 with another sample), non-European ancestry (>3 IQR from quartiles of European 1KGP samples on the first 4 PCs), or in sample exclusion lists (where available).

**Merged cohort QC.** Merge case and matched background control cohorts (using variants in common) and remove related samples (>.2 with another sample). Impute remaining genotypes and samples using TOPMed r2 panel.

**Post-imputation QC.** Remove imputed variants with low imputation quality ($R^2$<.3), low MAF (<.1%), and potential genotyping errors (genotyped variants with emperical $R^2$<.5 and imputed variants in LD - $r^2$>.5 with these variants).

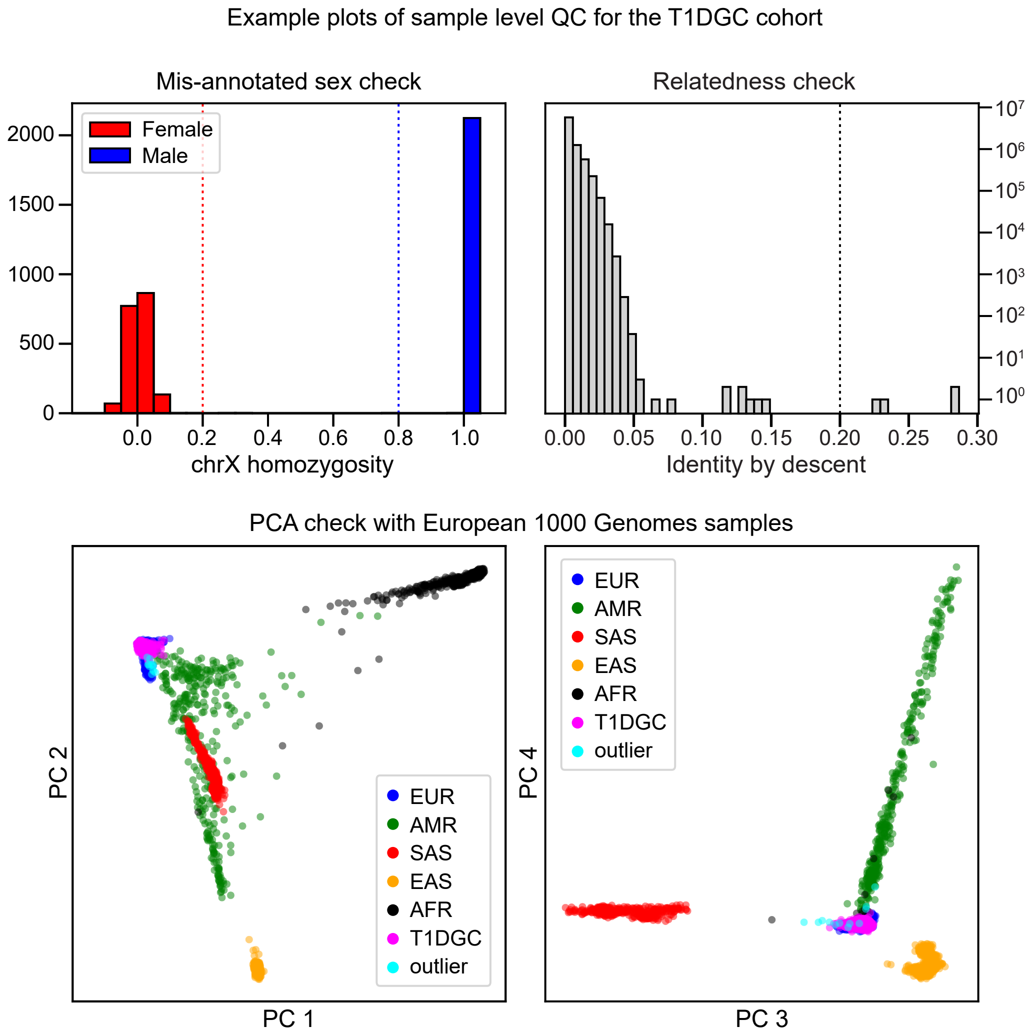


**Supplemental Figure 1. Flowchart of quality control steps for genotype imputation.** Flowchart (left) showing the steps of cohort-level, merged cohort, and post-imputation QC steps taken to filter out potential low quality variants and samples. Example plots (right) showing sample level QC steps for the T1DGC cohorts. Samples with ambiguous sex (dotted red and blue lines marks thresholds for females and males respectively), relatedness with another sample (dotted line marks threshold for relatedness), and non-European ancestry outliers (cyan points) are filtered out.

### Figure S2

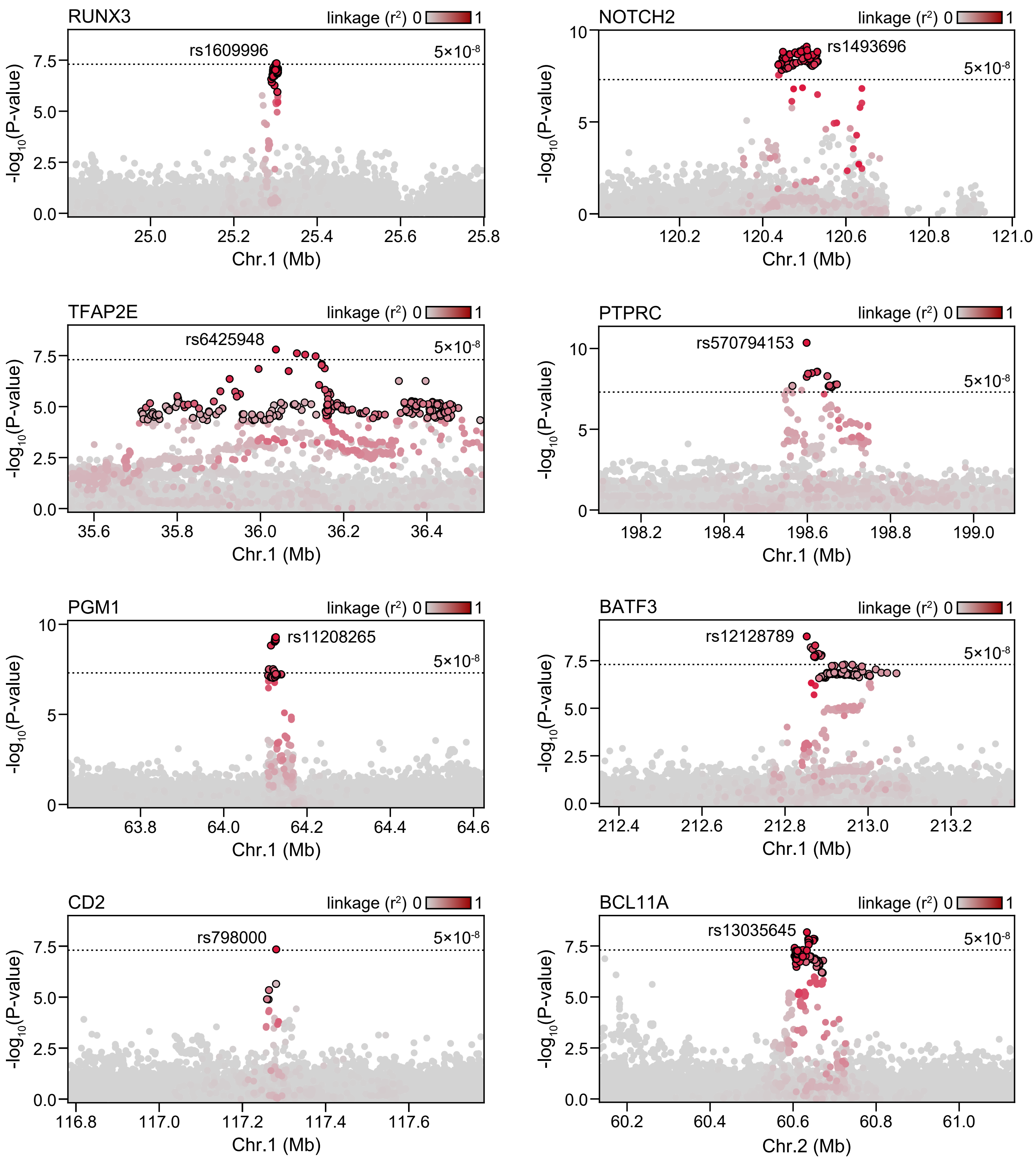
RUNX3
linkage (r²) 0 1
rs1609996
5×10⁻⁸
-log10(P-value)
Chr.1 (Mb)
NOTCH2
rs1493696
TFAP2E
rs6425948
PTPRC
rs570794153
PGM1
rs11208265
BATF3
rs12128789
CD2
rs798000
BCL11A
rs13035645
Chr.2 (Mb)

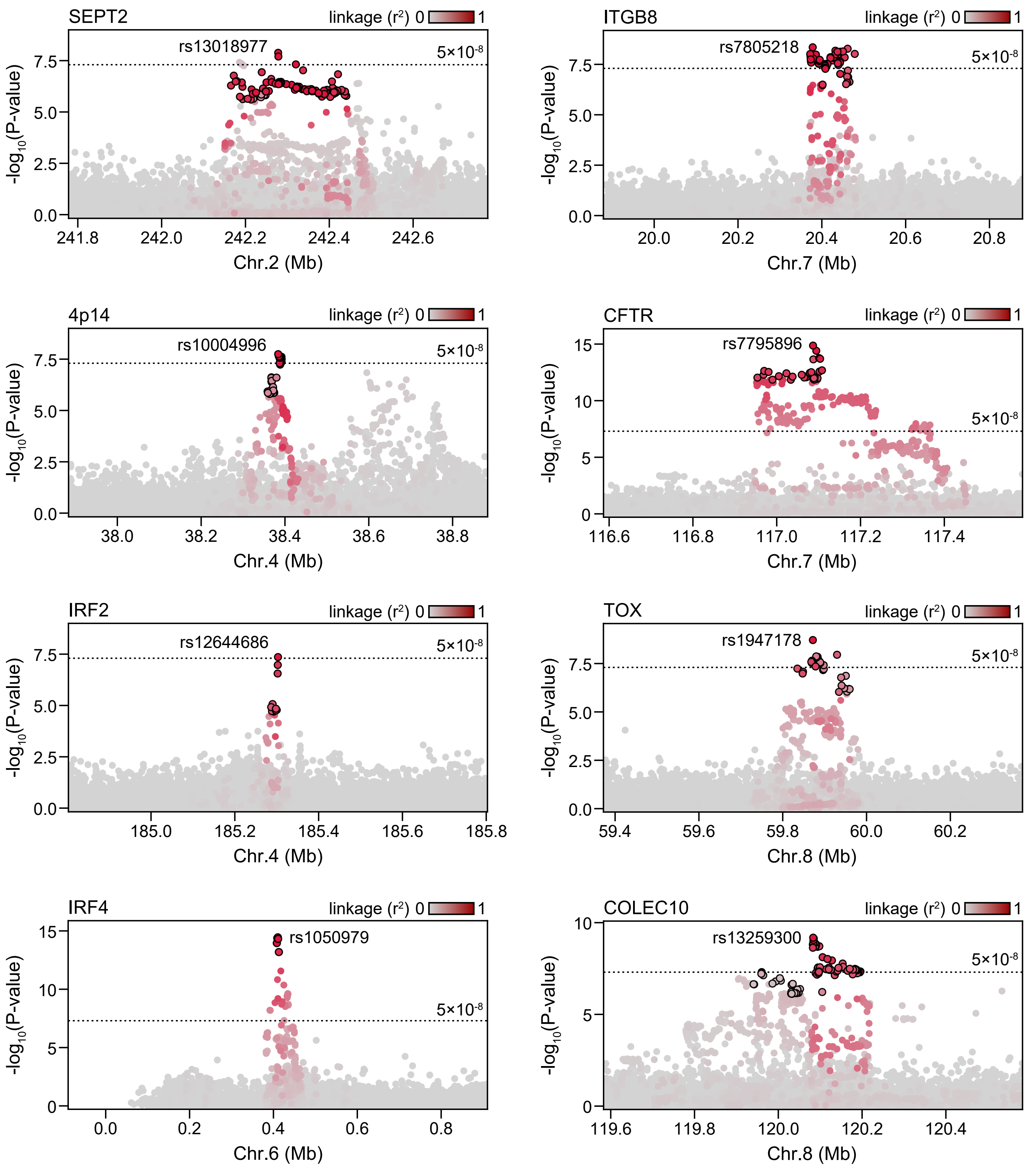

SEPT2
linkage (r2) 0 1
rs13018977
5×10-8
-log10(P-value)
Chr.2 (Mb)
ITGB8
rs7805218
Chr.7 (Mb)
4p14
rs10004996
Chr.4 (Mb)
CFTR
rs7795896
Chr.7 (Mb)
IRF2
rs12644686
Chr.4 (Mb)
TOX
rs1947178
Chr.8 (Mb)
IRF4
rs1050979
Chr.6 (Mb)
COLEC10
rs13259300
Chr.8 (Mb)

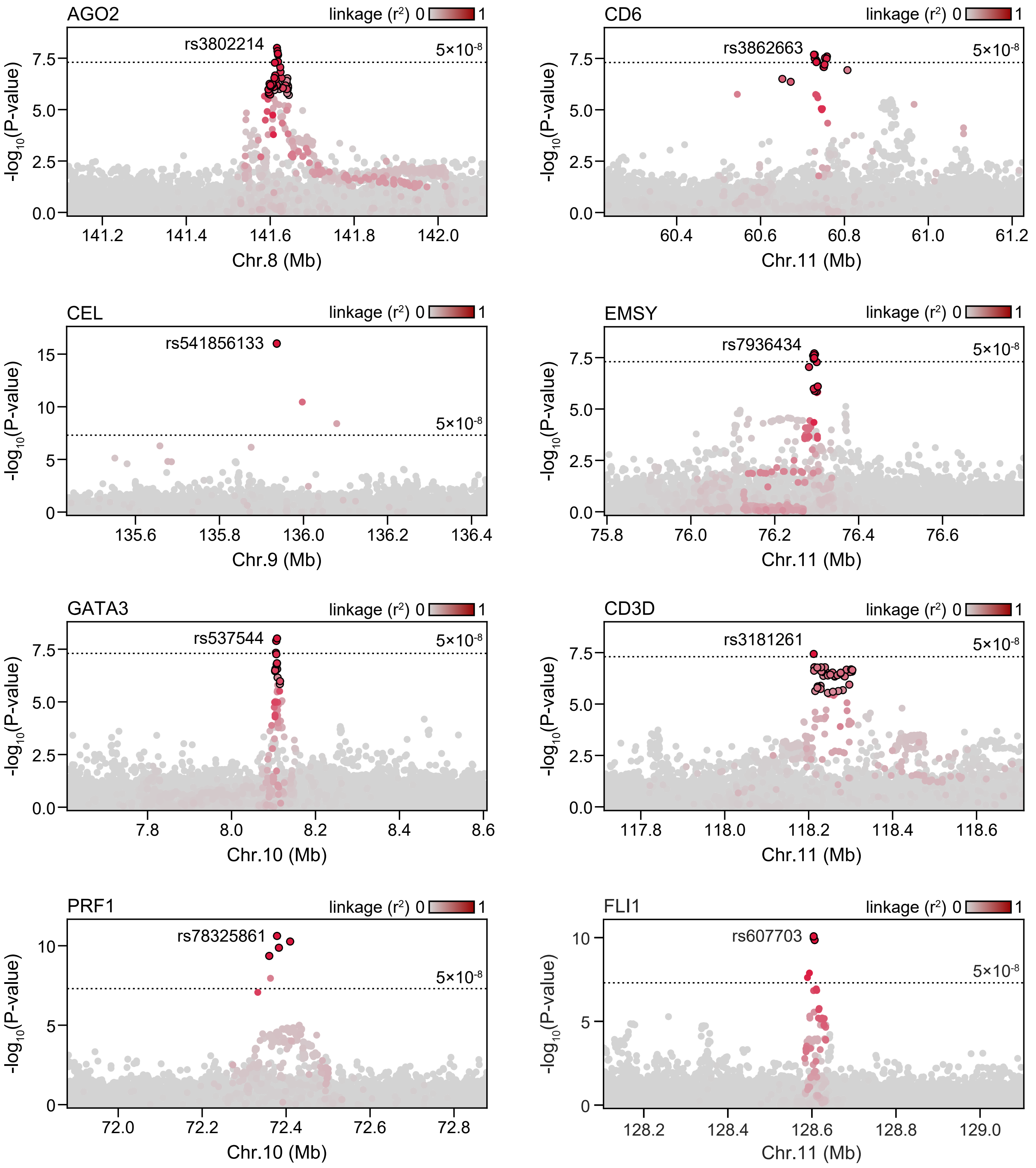
AGO2
linkage ($r^2$) 0 1
rs3802214
5×10$^{-8}$
$-log_{10}$(P-value)
0.0
2.5
5.0
7.5
141.2
141.4
141.6
141.8
142.0
Chr.8 (Mb)
CD6
linkage ($r^2$) 0 1
rs3862663
5×10$^{-8}$
$-log_{10}$(P-value)
0.0
2.5
5.0
7.5
60.4
60.6
60.8
61.0
61.2
Chr.11 (Mb)
CEL
linkage ($r^2$) 0 1
rs541856133
5×10$^{-8}$
$-log_{10}$(P-value)
0
5
10
15
135.6
135.8
136.0
136.2
136.4
Chr.9 (Mb)
EMSY
linkage ($r^2$) 0 1
rs7936434
5×10$^{-8}$
$-log_{10}$(P-value)
0.0
2.5
5.0
7.5
75.8
76.0
76.2
76.4
76.6
Chr.11 (Mb)
GATA3
linkage ($r^2$) 0 1
rs537544
5×10$^{-8}$
$-log_{10}$(P-value)
0.0
2.5
5.0
7.5
7.8
8.0
8.2
8.4
8.6
Chr.10 (Mb)
CD3D
linkage ($r^2$) 0 1
rs3181261
5×10$^{-8}$
$-log_{10}$(P-value)
0.0
2.5
5.0
7.5
117.8
118.0
118.2
118.4
118.6
Chr.11 (Mb)
PRF1
linkage ($r^2$) 0 1
rs78325861
5×10$^{-8}$
$-log_{10}$(P-value)
0
5
10
72.0
72.2
72.4
72.6
72.8
Chr.10 (Mb)
FLI1
linkage ($r^2$) 0 1
rs607703
5×10$^{-8}$
$-log_{10}$(P-value)
0
5
10
128.2
128.4
128.6
128.8
129.0
Chr.11 (Mb)

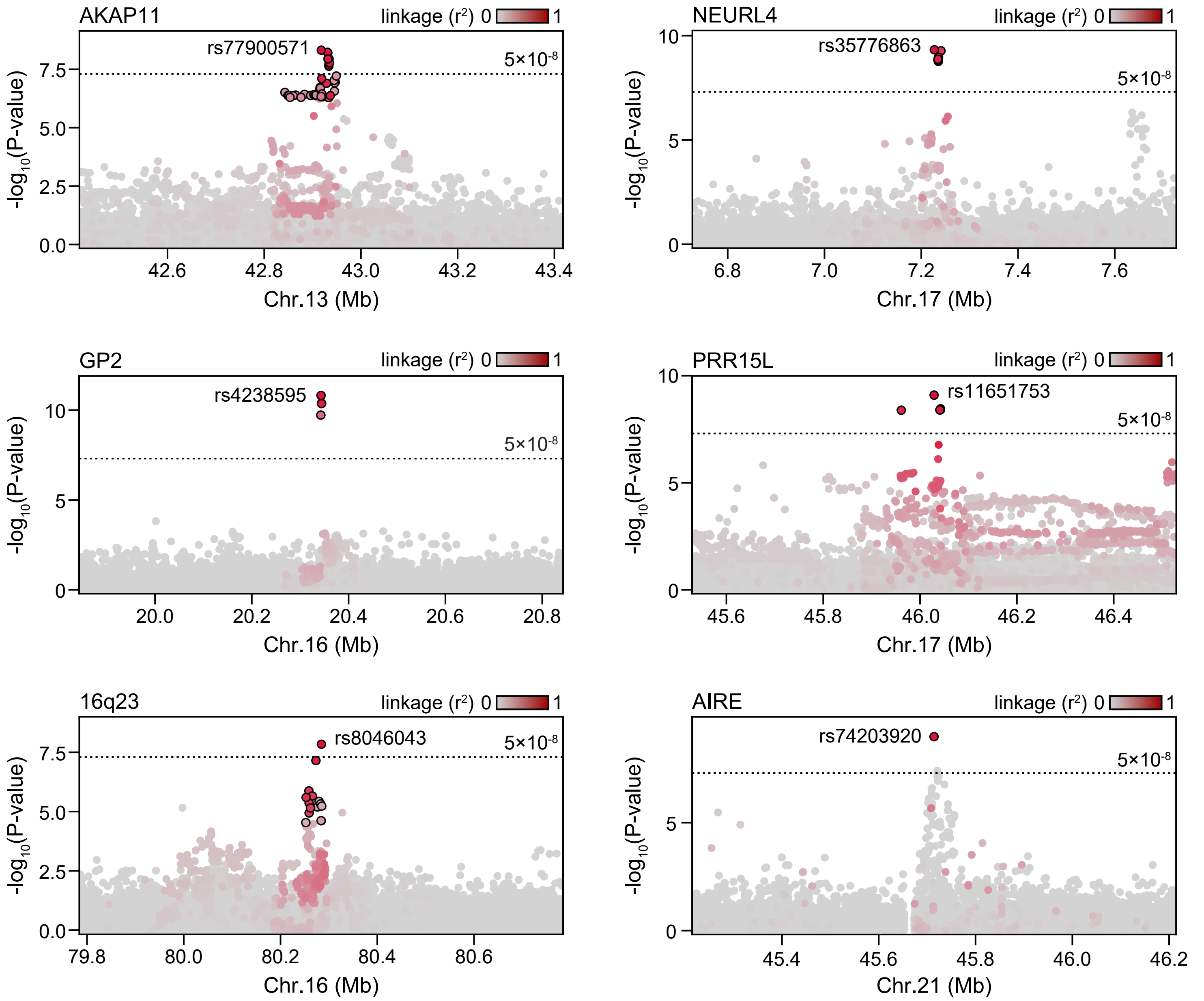


**Supplemental Figure 2. Locus plots of novel T1D loci.** Locus plots showing main signal association p-values for all variants in a 1 Mb window around the index variant for 30 novel T1D loci. Variants are colored based on linkage disequilibrium with the index variant. Circled variants are contained within the 99% credible set for the main signal at the locus.

### Figure S3

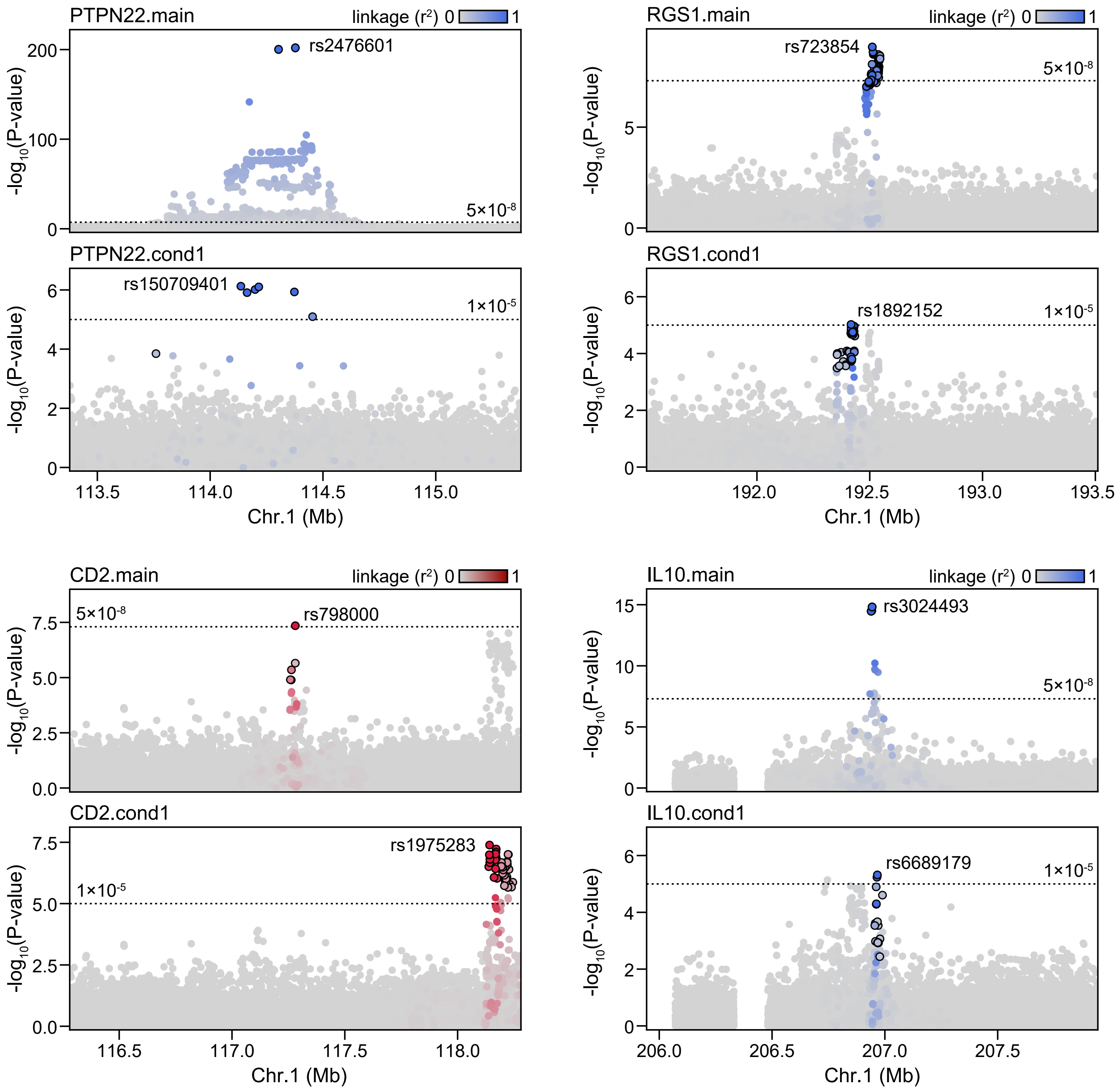

PTPN22.main
linkage (r²) 0 1
rs2476601
5×10-8
-log10(P-value)
PTPN22.cond1
rs150709401
1×10-5
113.5
114.0
114.5
115.0
Chr.1 (Mb)
RGS1.main
rs723854
RGS1.cond1
rs1892152
192.0
192.5
193.0
193.5
CD2.main
rs798000
CD2.cond1
rs1975283
116.5
117.0
117.5
118.0
IL10.main
rs3024493
IL10.cond1
rs6689179
206.0
206.5
207.0
207.5

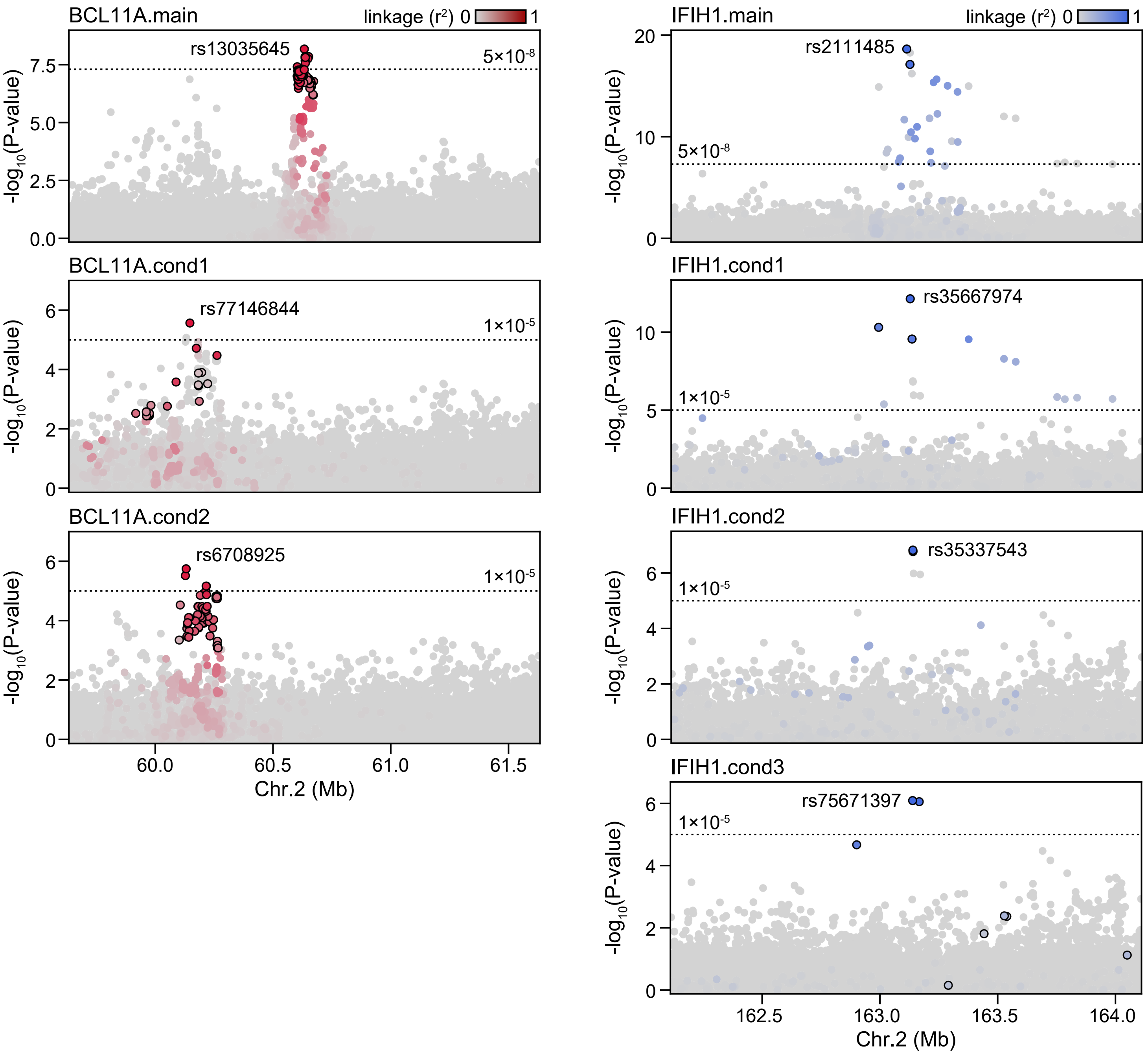

BCL11A.main
linkage (r²) 0 1
rs13035645
5×10-8
-log10(P-value)
BCL11A.cond1
rs77146844
1×10-5
BCL11A.cond2
rs6708925
Chr.2 (Mb)
IFIH1.main
rs2111485
IFIH1.cond1
rs35667974
IFIH1.cond2
rs35337543
IFIH1.cond3
rs75671397

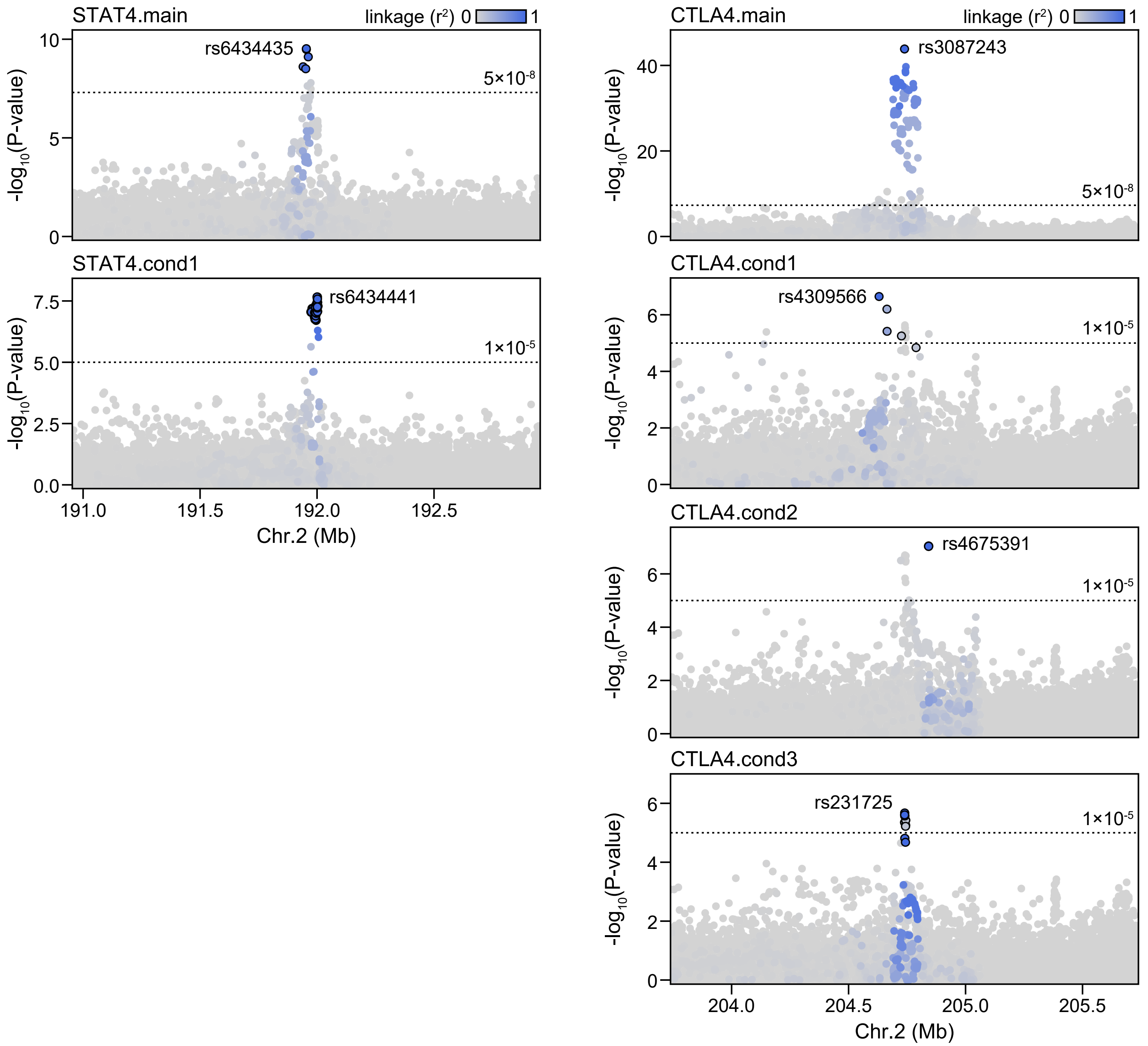

STAT4.main
linkage (r2) 0 1
rs6434435
5×10-8
-log10(P-value)
STAT4.cond1
rs6434441
1×10-5
191.0
191.5
192.0
192.5
Chr.2 (Mb)
CTLA4.main
rs3087243
CTLA4.cond1
rs4309566
CTLA4.cond2
rs4675391
CTLA4.cond3
rs231725
204.0
204.5
205.0
205.5
Chr.2 (Mb)

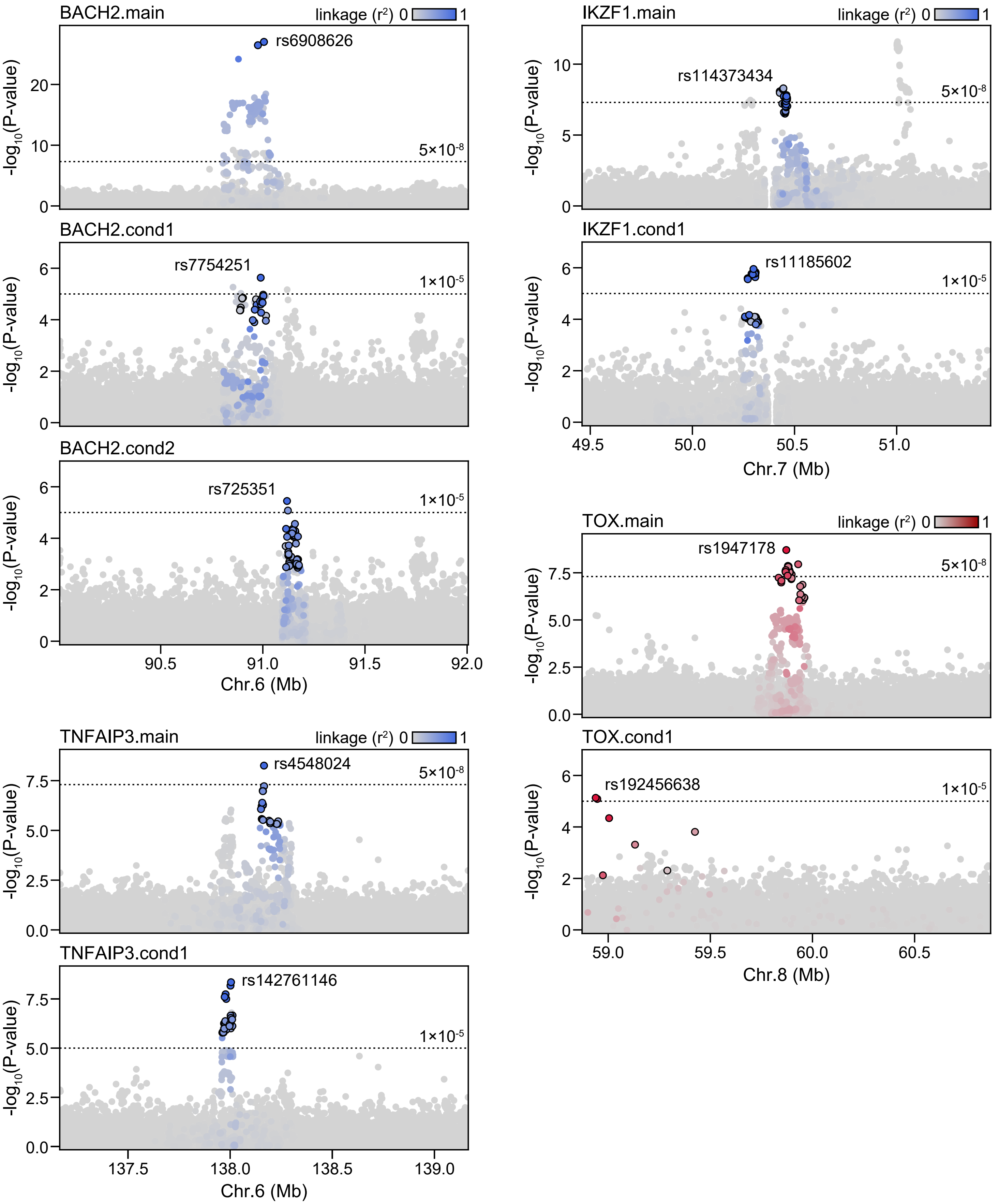
BACH2.main
linkage (r²) 0 1
rs6908626
5×10⁻⁸
-log10(P-value)
BACH2.cond1
rs7754251
1×10⁻⁵
BACH2.cond2
rs725351
1×10⁻⁵
90.5
91.0
91.5
92.0
Chr.6 (Mb)
IKZF1.main
rs114373434
5×10⁻⁸
IKZF1.cond1
rs11185602
1×10⁻⁵
49.5
50.0
50.5
51.0
Chr.7 (Mb)
TOX.main
rs1947178
5×10⁻⁸
TOX.cond1
rs192456638
1×10⁻⁵
59.0
59.5
60.0
60.5
Chr.8 (Mb)
TNFAIP3.main
rs4548024
5×10⁻⁸
TNFAIP3.cond1
rs142761146
1×10⁻⁵
137.5
138.0
138.5
139.0
Chr.6 (Mb)

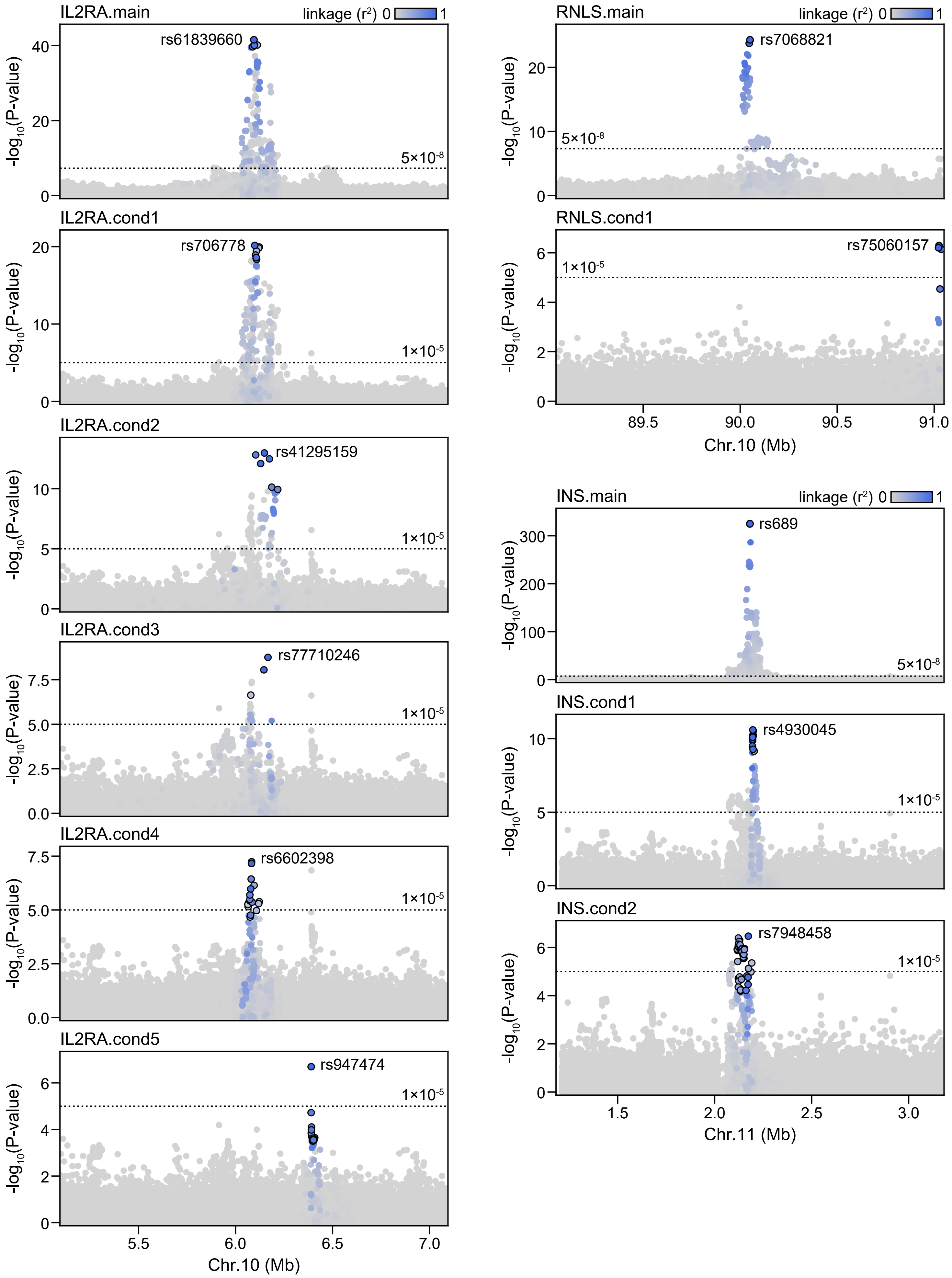
IL2RA.main
linkage (r²) 0 1
rs61839660
5×10⁻⁸
IL2RA.cond1
rs706778
1×10⁻⁵
IL2RA.cond2
rs41295159
1×10⁻⁵
IL2RA.cond3
rs77710246
1×10⁻⁵
IL2RA.cond4
rs6602398
1×10⁻⁵
IL2RA.cond5
rs947474
1×10⁻⁵
-log10(P-value)
5.5
6.0
6.5
7.0
Chr.10 (Mb)
RNLS.main
linkage (r²) 0 1
rs7068821
5×10⁻⁸
RNLS.cond1
rs75060157
1×10⁻⁵
89.5
90.0
90.5
91.0
Chr.10 (Mb)
INS.main
linkage (r²) 0 1
rs689
5×10⁻⁸
INS.cond1
rs4930045
1×10⁻⁵
INS.cond2
rs7948458
1×10⁻⁵
1.5
2.0
2.5
3.0
Chr.11 (Mb)

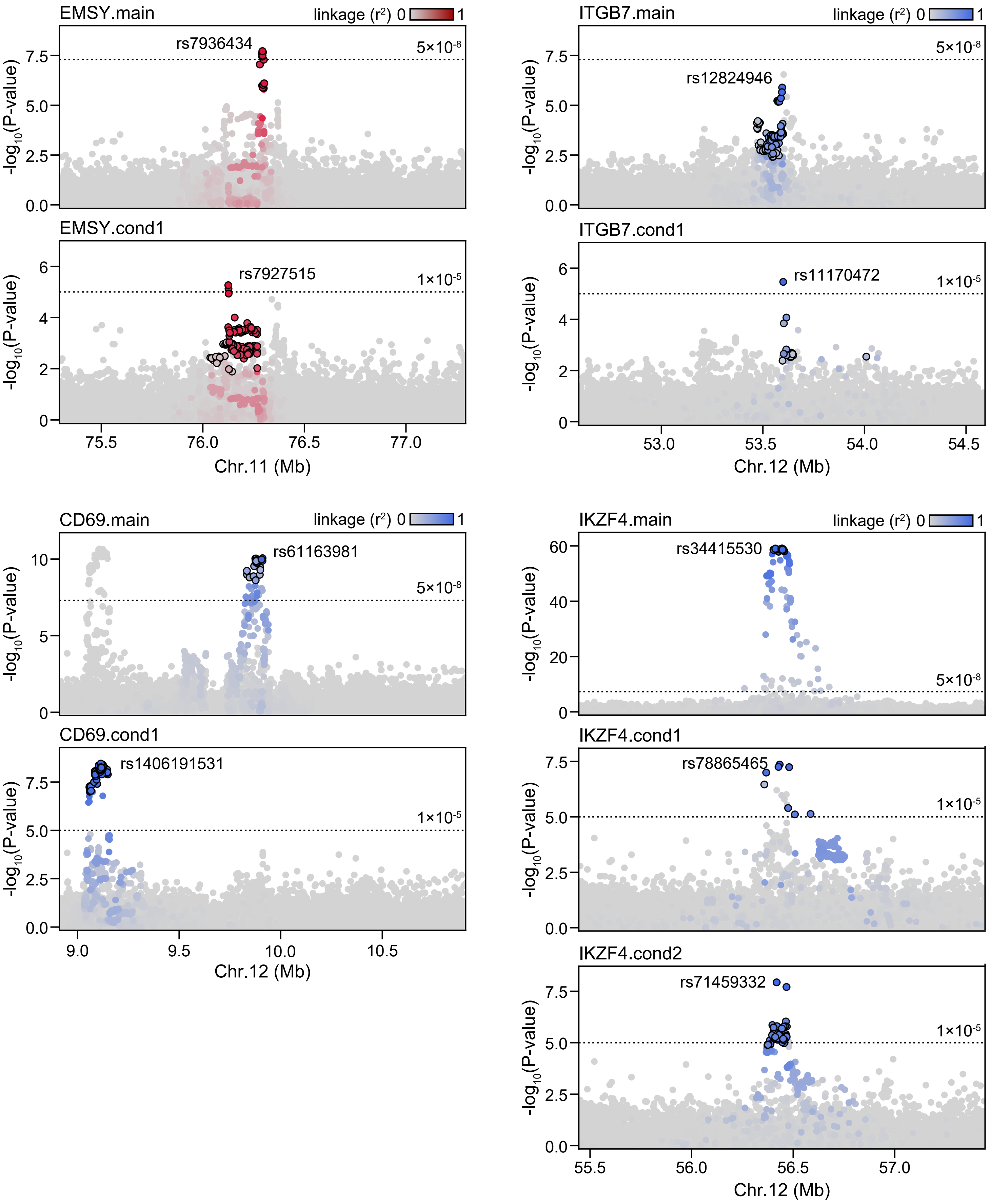

EMSY.main
linkage (r²) 0 1
rs7936434
5×10⁻⁸
-log10(P-value)
7.5
5.0
2.5
0.0
EMSY.cond1
rs7927515
1×10⁻⁵
6
4
2
0
75.5
76.0
76.5
77.0
Chr.11 (Mb)
ITGB7.main
rs12824946
ITGB7.cond1
rs11170472
53.0
53.5
54.0
54.5
Chr.12 (Mb)
CD69.main
rs61163981
10
5
CD69.cond1
rs1406191531
9.0
9.5
10.0
10.5
IKZF4.main
rs34415530
60
40
20
IKZF4.cond1
rs78865465
IKZF4.cond2
rs71459332
55.5
56.0
56.5
57.0

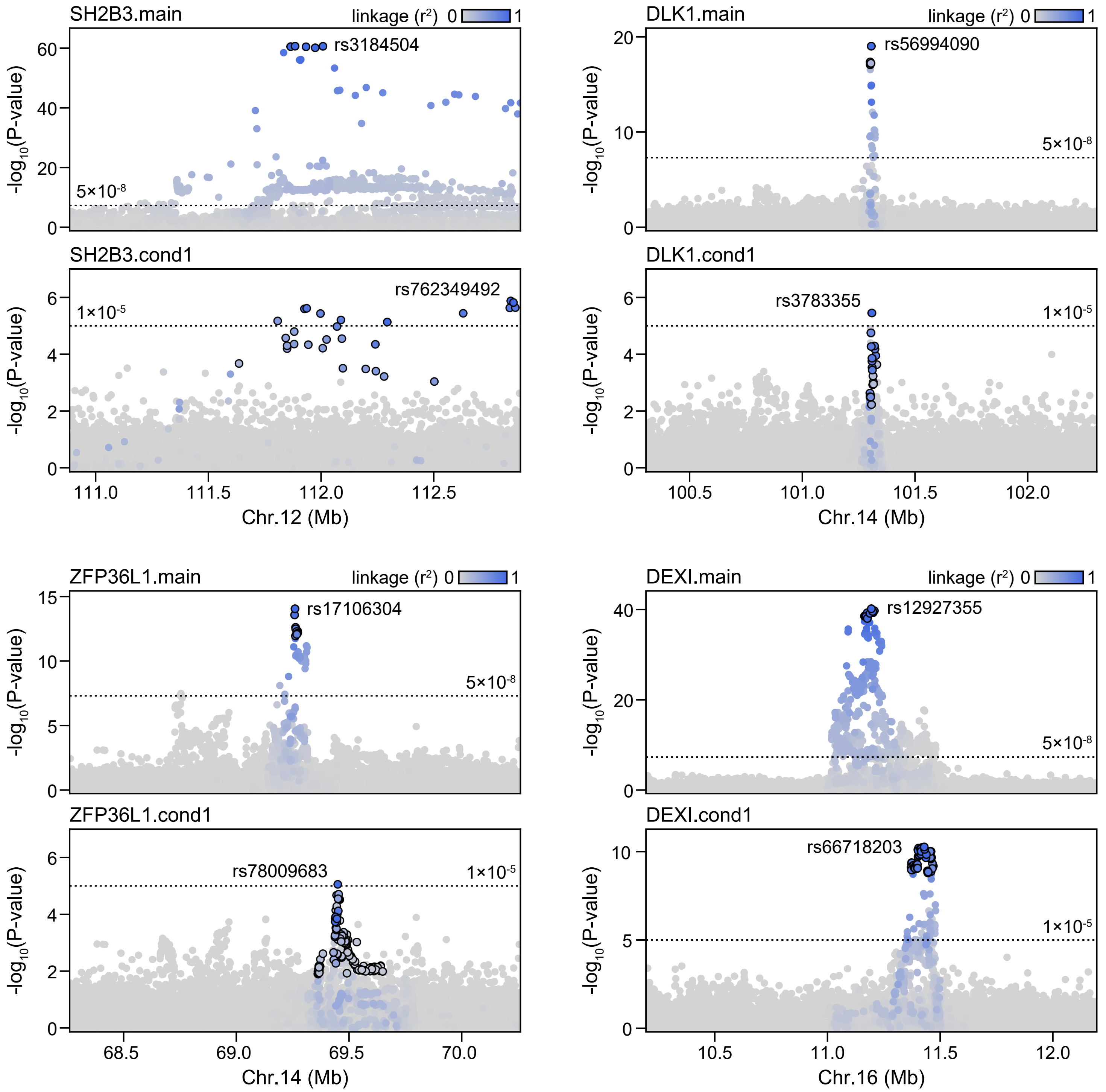

SH2B3.main
linkage (r²) 0 1
rs3184504
5×10⁻⁸
SH2B3.cond1
rs762349492
1×10⁻⁵
-log10(P-value)
111.0
111.5
112.0
112.5
Chr.12 (Mb)
DLK1.main
rs56994090
5×10⁻⁸
DLK1.cond1
rs3783355
1×10⁻⁵
100.5
101.0
101.5
102.0
Chr.14 (Mb)
ZFP36L1.main
rs17106304
5×10⁻⁸
ZFP36L1.cond1
rs78009683
1×10⁻⁵
68.5
69.0
69.5
70.0
Chr.14 (Mb)
DEXI.main
rs12927355
5×10⁻⁸
DEXI.cond1
rs66718203
1×10⁻⁵
10.5
11.0
11.5
12.0
Chr.16 (Mb)

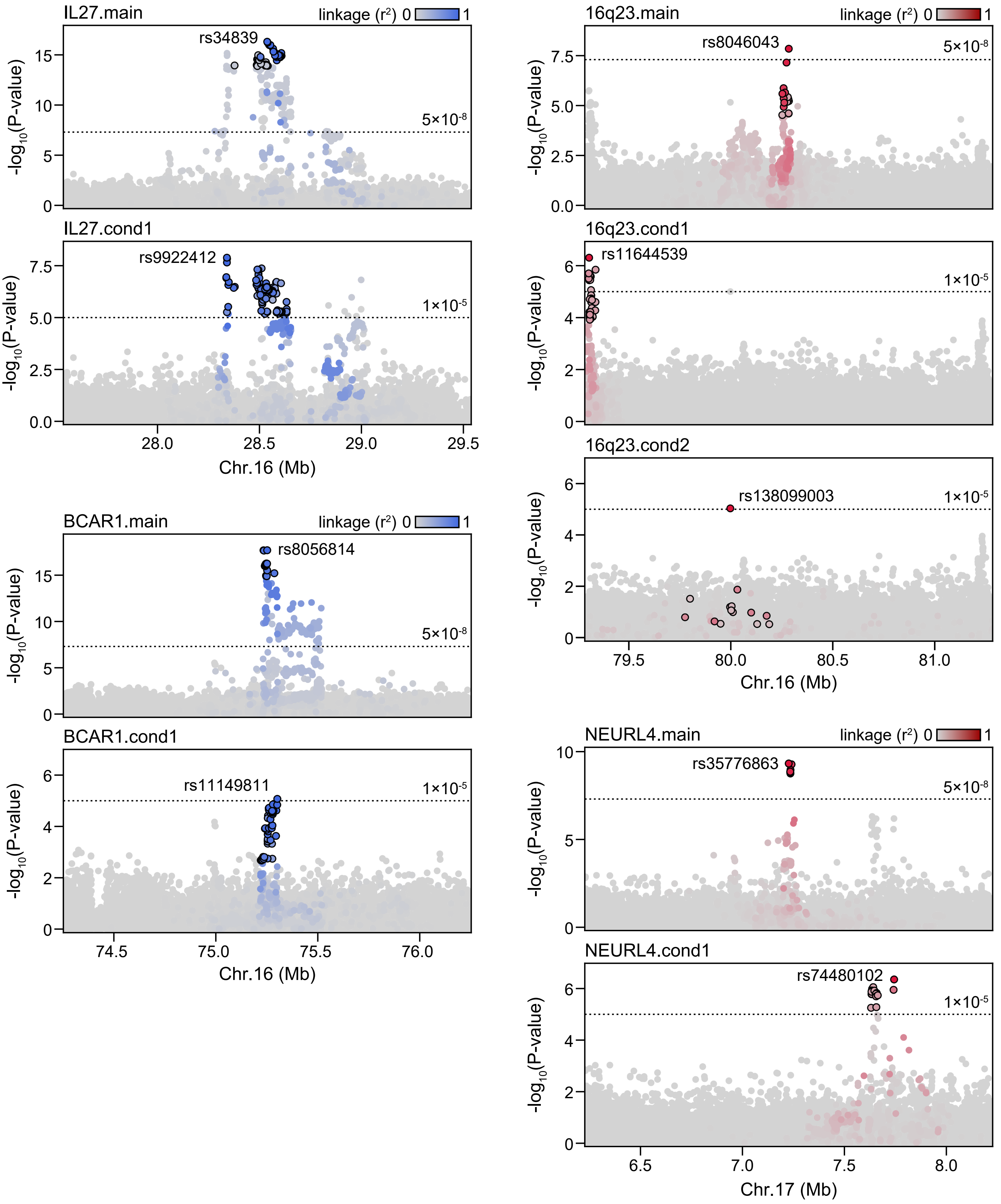

IL27.main
linkage (r²) 0 1
rs34839
5×10-8
-log10(P-value)
IL27.cond1
rs9922412
1×10-5
28.0
28.5
29.0
29.5
Chr.16 (Mb)
BCAR1.main
rs8056814
BCAR1.cond1
rs11149811
74.5
75.0
75.5
76.0
16q23.main
rs8046043
16q23.cond1
rs11644539
16q23.cond2
rs138099003
79.5
80.0
80.5
81.0
NEURL4.main
rs35776863
NEURL4.cond1
rs74480102
6.5
7.0
7.5
8.0
Chr.17 (Mb)

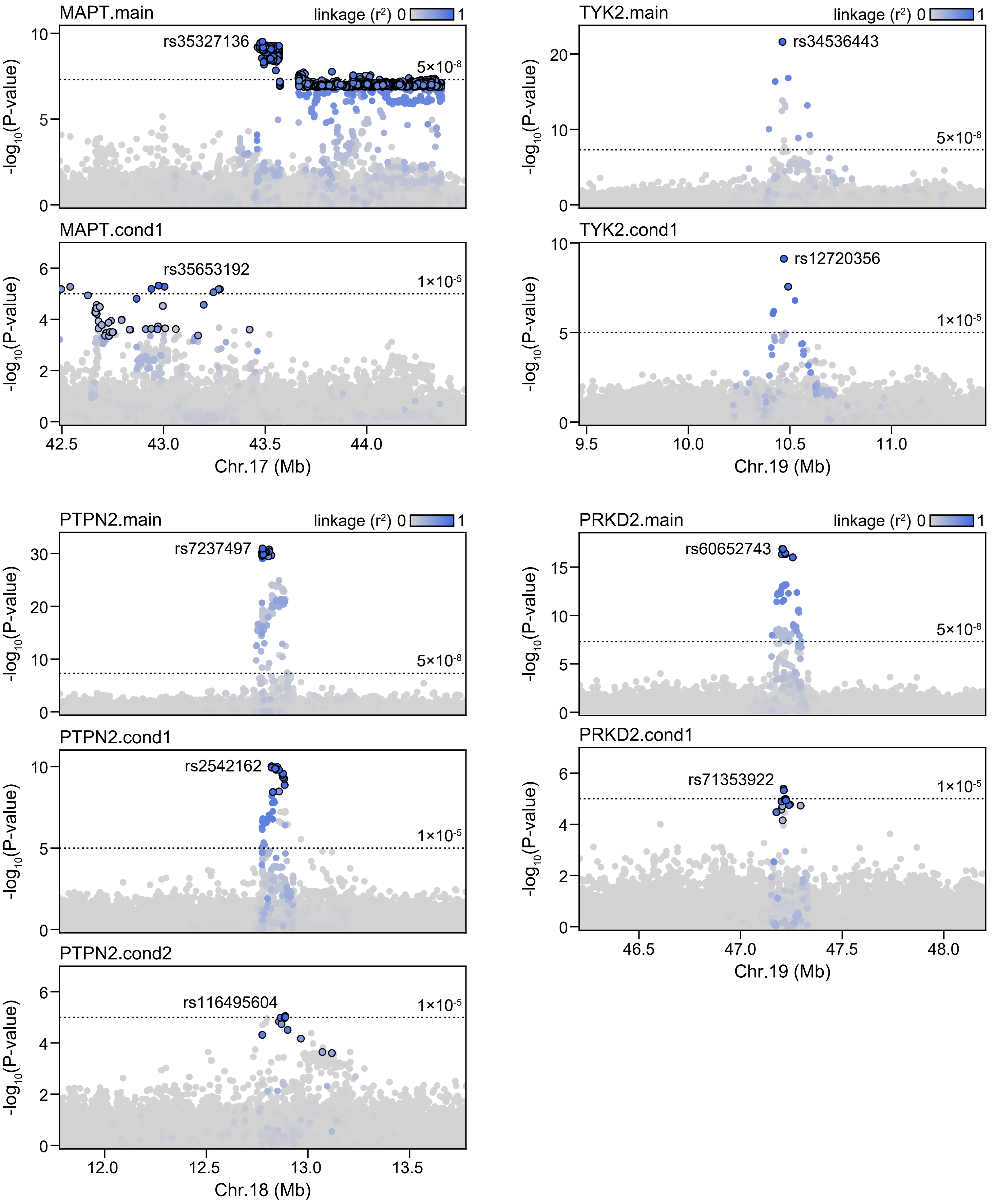
MAPT.main
linkage ($r^2$) 0 1
rs35327136
$5\times10^{-8}$
MAPT.cond1
rs35653192
$1\times10^{-5}$
42.5
43.0
43.5
44.0
Chr.17 (Mb)
TYK2.main
linkage ($r^2$) 0 1
rs34536443
$5\times10^{-8}$
TYK2.cond1
rs12720356
$1\times10^{-5}$
9.5
10.0
10.5
11.0
Chr.19 (Mb)
PTPN2.main
linkage ($r^2$) 0 1
rs7237497
$5\times10^{-8}$
PTPN2.cond1
rs2542162
$1\times10^{-5}$
PTPN2.cond2
rs116495604
$1\times10^{-5}$
12.0
12.5
13.0
13.5
Chr.18 (Mb)
PRKD2.main
linkage ($r^2$) 0 1
rs60652743
$5\times10^{-8}$
PRKD2.cond1
rs71353922
$1\times10^{-5}$
46.5
47.0
47.5
48.0
Chr.19 (Mb)
$-\log_{10}$(P-value)

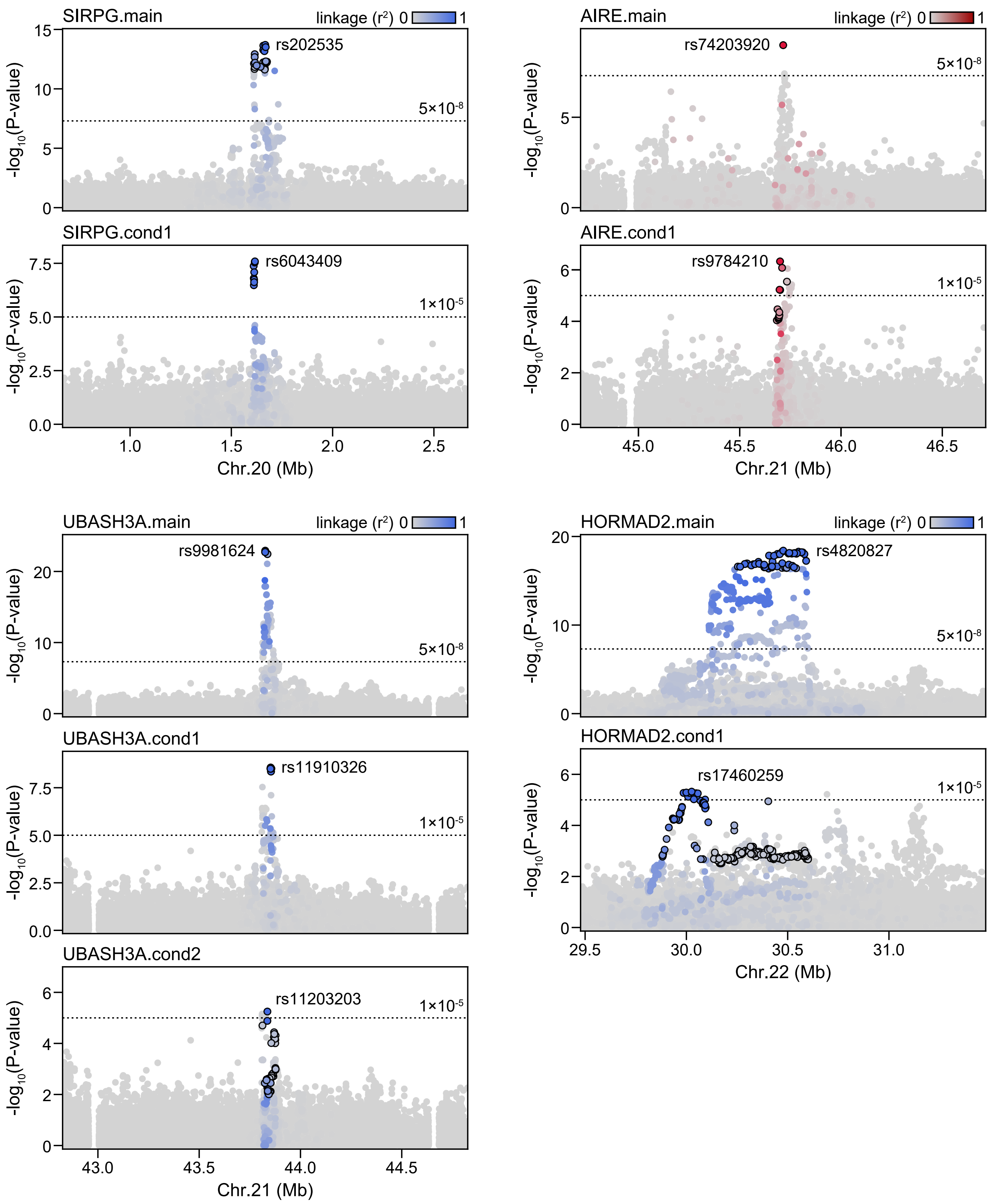

SIRPG.main
linkage (r²) 0 1
rs202535
5×10-8
-log10(P-value)
SIRPG.cond1
rs6043409
1×10-5
Chr.20 (Mb)
AIRE.main
rs74203920
AIRE.cond1
rs9784210
Chr.21 (Mb)
UBASH3A.main
rs9981624
UBASH3A.cond1
rs11910326
UBASH3A.cond2
rs11203203
Chr.21 (Mb)
HORMAD2.main
rs4820827
HORMAD2.cond1
rs17460259
Chr.22 (Mb)

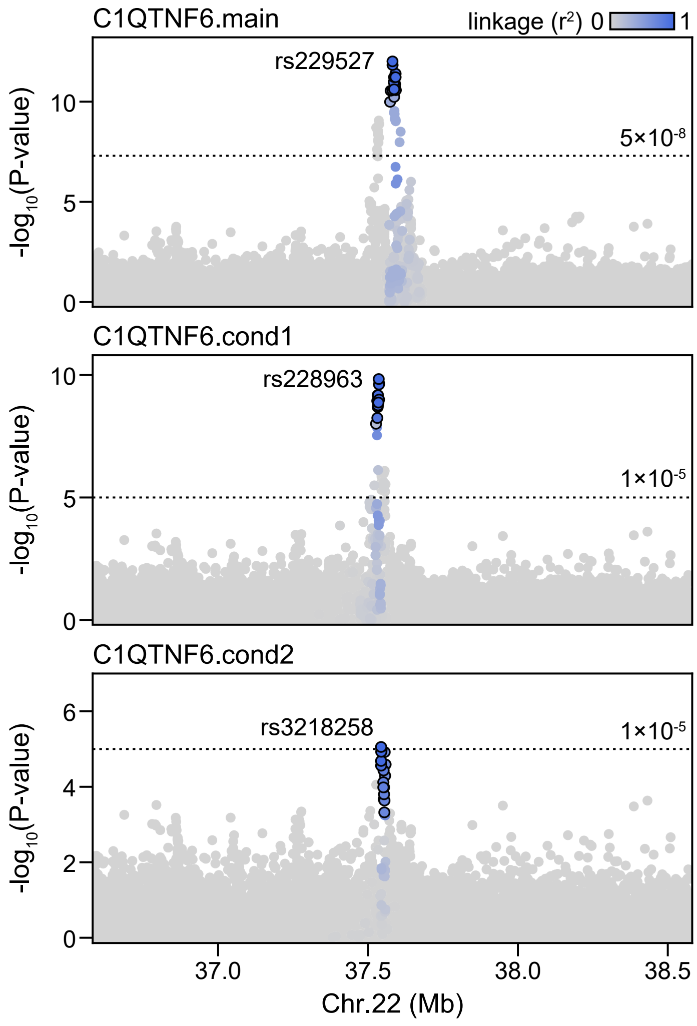


**Supplemental Figure 3. Locus plots of conditional T1D signals.** Locus plots showing main and conditional association signals for loci with multiple independent signals. Variants in 2 Mb windows around the index variant (labeled) are colored by linkage disequilibrium ($r^2$) with the index variant (blue for known loci and red for novel loci) and are circled if they are included in the 99% credible set for each signal.

### Figure S4

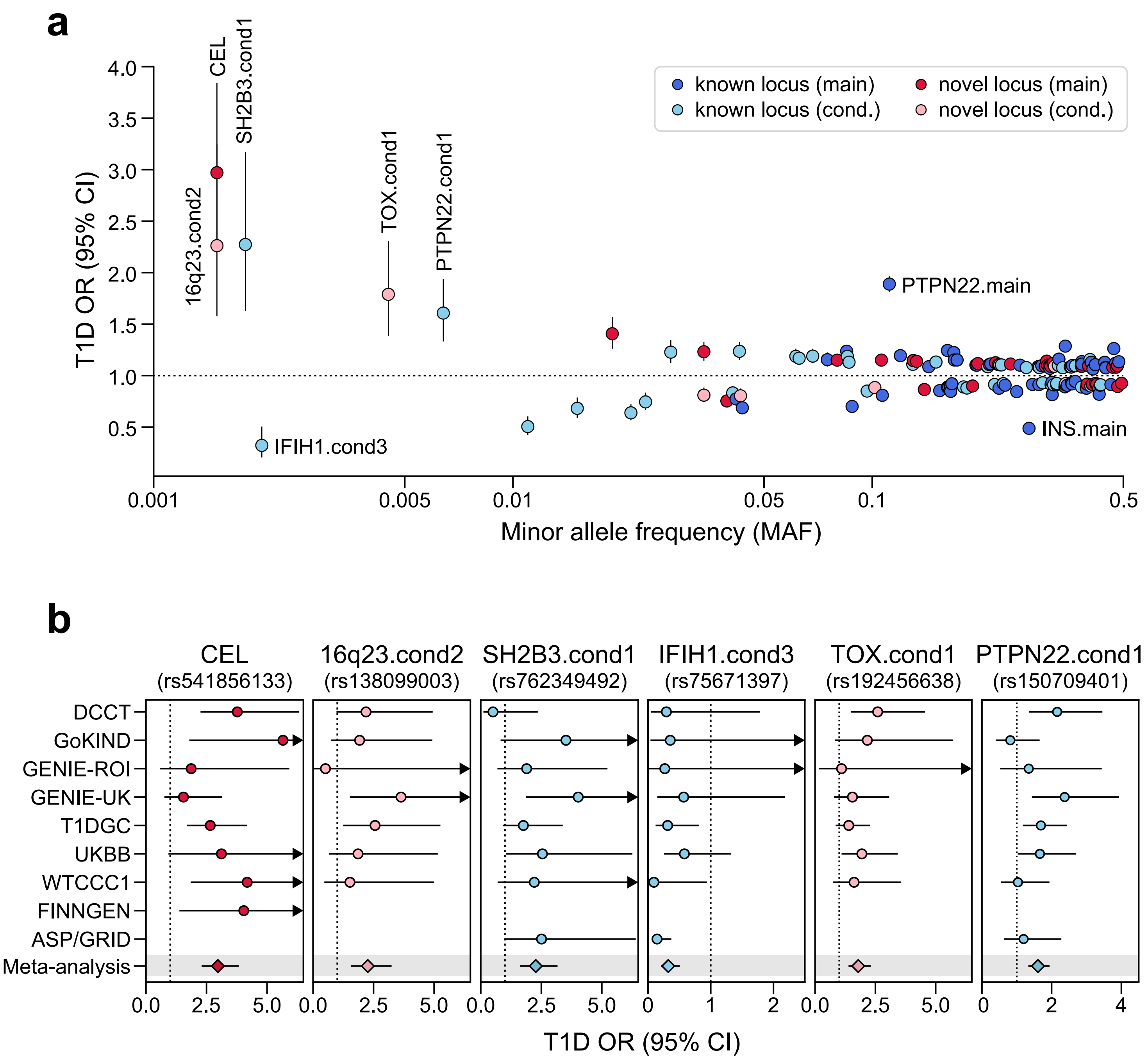


**Supplemental Figure 4. Rare variants with large effects on T1D risk.** (a) The relationship between minor allele frequency and T1D odds ratios (OR) for 141 non-MHC risk signals. Points represent GWAS OR estimates and lines represent 95% confidence intervals. (b) Comparison of odds ratios across cohorts for rare variants with MAF<0.01. For conditional signals, odds ratio estimates are from conditional analyses. Missing values indicate that the variant was not tested in the cohort.

### Figure S5

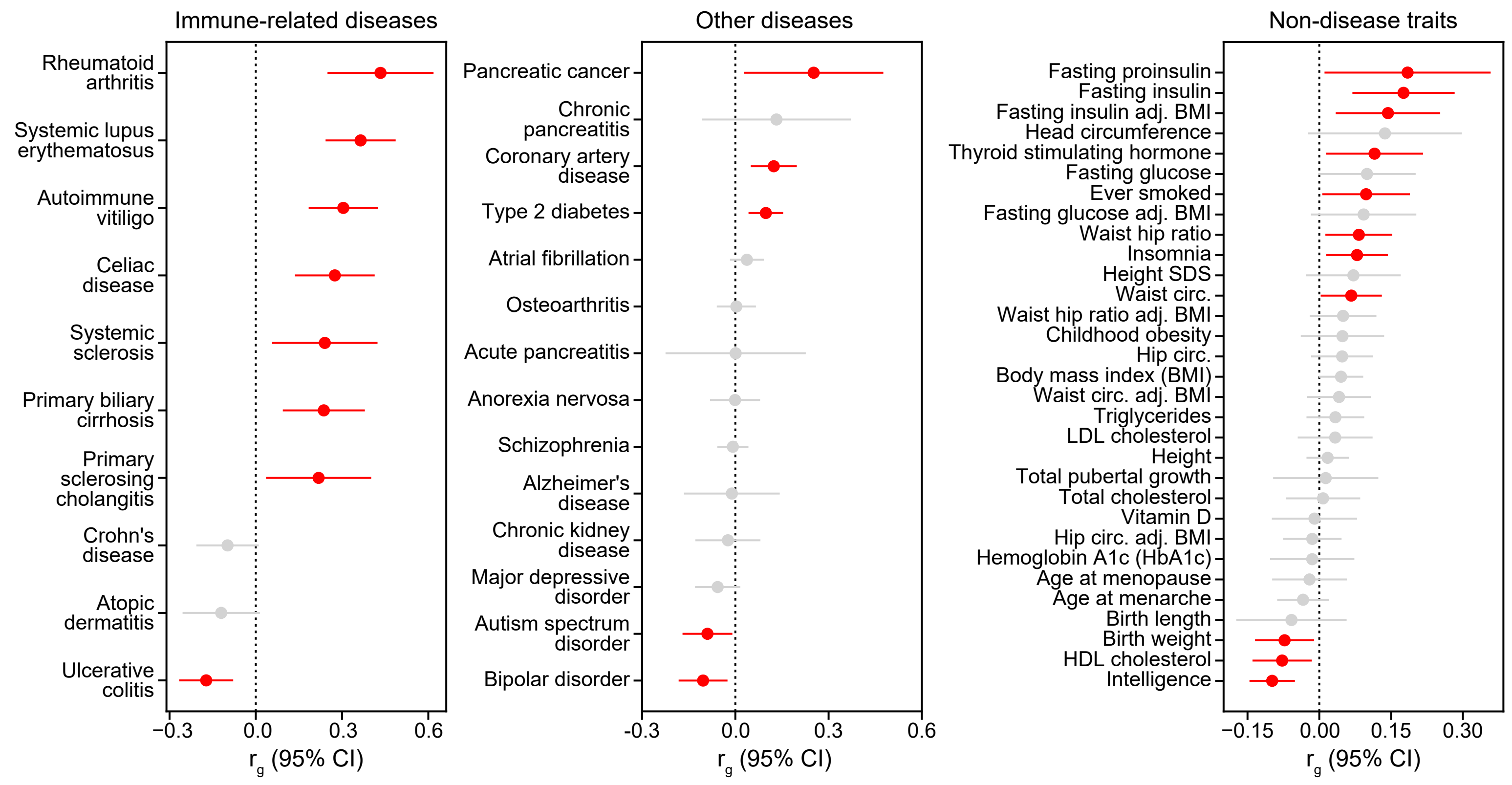


**Supplemental Figure 5. Type 1 diabetes genetic correlations.** Genetic correlations between T1D and other traits, including immune-related diseases (left), other diseases (middle), and non-disease traits (right), adj.=adjusted, circ.=circumference. Points represent genetic correlation estimates and lines represent 95% confidence intervals. Colors indicate significance: red indicates that the correlation is significant after FDR correction (FDR<0.1) and grey indicates that the correlation is not significant.

### Figure S6

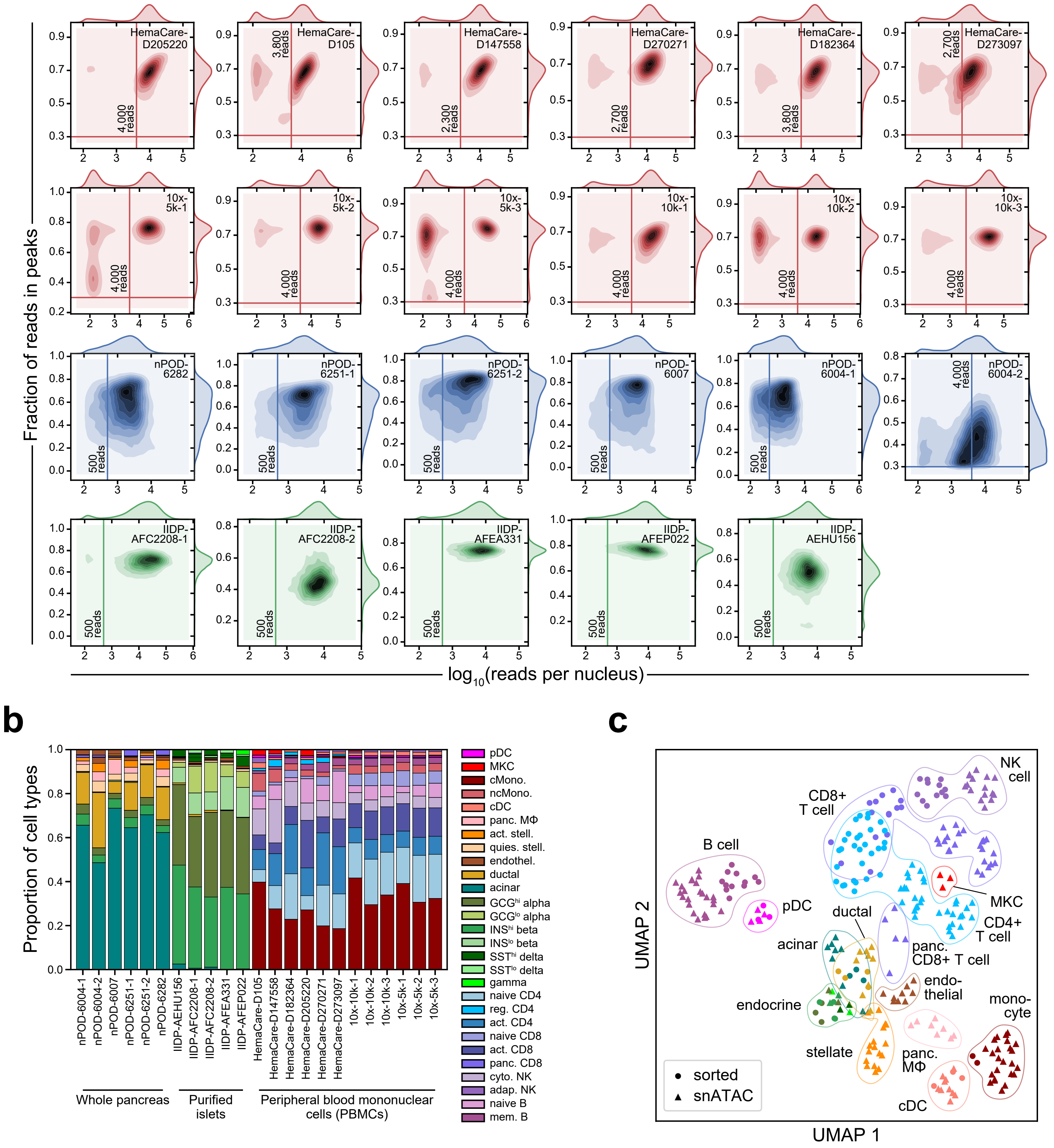


**Supplemental Figure 6. snATAC-seq quality control metrics and comparison to sorted datasets.** (a) Kernel density of the number of unique reads (log-transformed) compared to the fraction of reads in bulk peaks per cell. Thresholds for unique reads range between 500-4000 as indicated on each plot, and an additional threshold of fraction of reads in peaks > 0.3 was set for datasets generated using 10x Genomics technology. (b) Proportion of cell types within each experiment for cells passing QC. (c) Comparison of pseudobulk snATAC-seq profiles (within donor, at least 50 cells) to FACS purified ATAC-seq for major pancreatic or unstimulated immune cell types using PCA+ UMAP dimensionality reduction on read counts within 448,142 merged cREs from snATAC-seq clusters.

### Figure S7

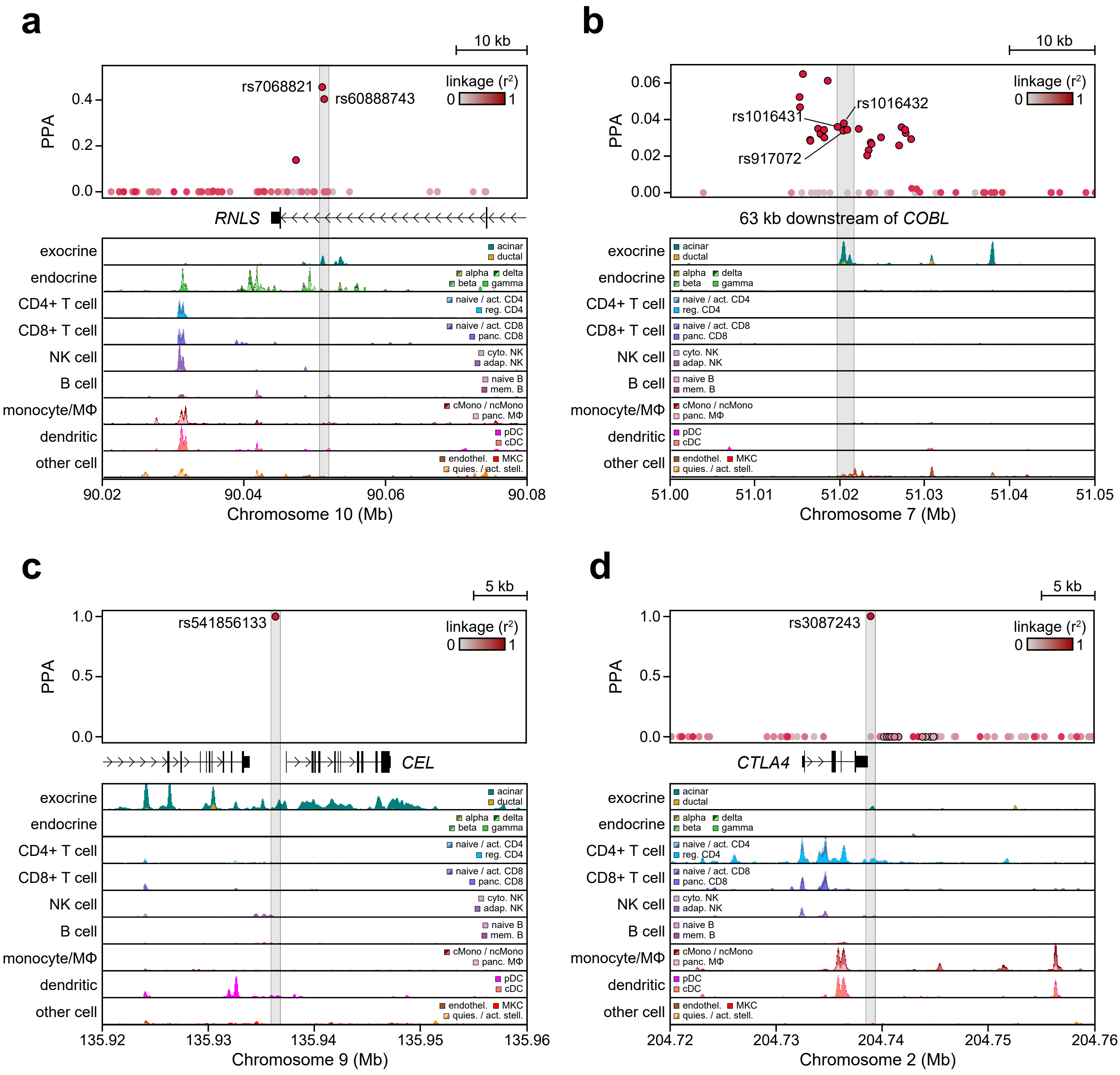


**Supplemental Figure 7. Fine mapped variants in acinar regulatory elements.** (a) The *RNLS* (main) signal contains two variants (rs7068821, PPA=0.455; rs60888743, PPA=0.403) located in an acinar-specific intronic peak within *RNLS* (chr10:90,020,000-90,080,000, scale: 0-10 CPM). (b) The *COBL* signal contains three variants (rs917072, PPA=0.034; rs1016432, PPA=0.038; rs1016431, PPA=0.034) located in an acinar-specific distal peak 63 kb downstream of *COBL* (chr7:51,000,000-51,050,000, scale: 0-20 CPM). (c) The *CEL* signal contains a single fine mapped rare variant (rs541856133, PPA=1.0) in an acinar-specific broad region of chromatin accessibility directly upstream of the *CEL* promoter (chr9:135,920,000-135,960,000, scale: 0-5 CPM). (d) The *CTLA4* (main) signal contains a single fine mapped variant (rs3087243, PPA=1.0) downstream of CTLA4 in an acinar peak and broad region of chromatin accessibility in regulatory T cells (chr2:204,720,000-204,760,000, scale: 0-5 CPM).

### Figure S8

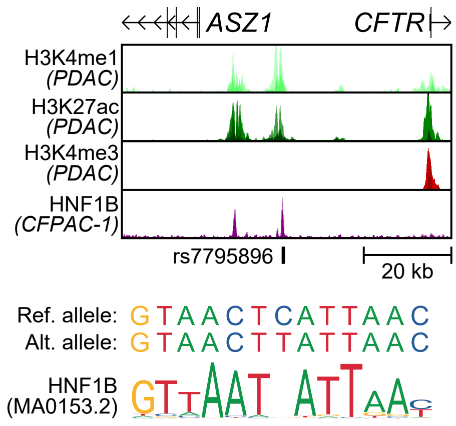


**Supplemental Figure 8. rs7795896 overlaps ChIP-seq peaks for HNF1B in ductal cell models.** Genome browser (region: chr7:117,050,000-117,125,000) showing rs7795896 overlaps peaks for histone marks of active enhancers (H3K4me1, scale: 0-30; H3K27ac, scale: 0-100) but not promoters (H3K4me3, scale: 0-300) in pancreatic ductal adenocarcinoma cell lines (PDAC: Capan-1, Capan-2, and CFPAC-1). rs7795896 also overlaps a ChIP-seq peak for the transcription factor HNF1B (scale: 0-30) in CFPAC-1 cells and a predicted HNF1B sequence motif.

### Figure S9

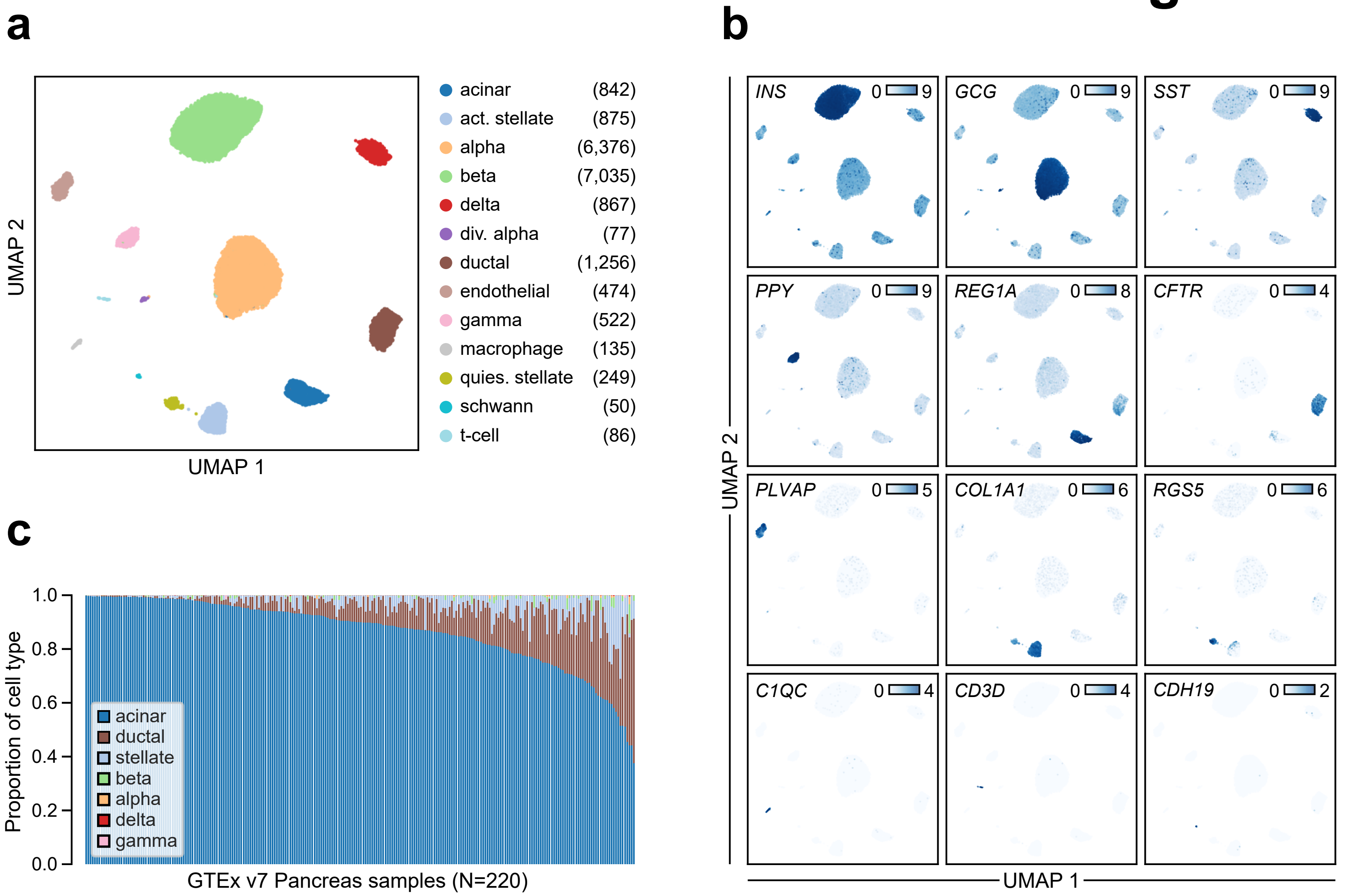


**Supplemental Figure 9. Islet single cell RNA-seq clusters and estimation of pancreas cell type proportions.** (a) Re-analysis of an islet single cell RNA-seq dataset consisting of 12 donors finds 13 clusters comprised of 18,844 total cells, plotted on UMAP coordinates and colored by cluster assignment. The number of cells within each cluster is indicated in parentheses to the right of the cluster label. div. alpha: dividing alpha, act. stellate: activated stellate, quies. stellate: quiescent stellate. (b) Single cell expression (in log-transformed counts) of key marker genes for each cluster plotted on UMAP coordinates. (c) Cell type proportions estimated by MuSiC for 220 bulk pancreas RNA-seq samples from the GTEx v7 release. Stellate cell proportions are combined estimates for activated and quiescent stellate cells.
